# Friedreich ataxia transcriptomic dysregulation and identification of cell type-specific biomarkers: A systematic review and meta-analysis

**DOI:** 10.64898/2026.03.18.712785

**Authors:** Marnie L. Maddock, Sara Miellet, Anjila Dongol, Amy J. Hulme, Chloe K. Kennedy, Louise A. Corben, Rocio K. Finol-Urdaneta, Alberto Nettel-Aguirre, Chiara Dionisi, Martin B. Delatycki, Joel M. Gottesfeld, Massimo Pandolfo, Elisabetta Soragni, Sanjay I. Bidichandani, Jarmon G. Lees, Shiang Y. Lim, Jill S. Napierala, Marek Napierala, Mirella Dottori

## Abstract

Friedreich ataxia (FRDA) is a progressive multisystem neurodegenerative disease mostly caused by a homozygous GAA repeat expansion in the *FXN* gene, leading to deficiency of the protein frataxin. Despite ubiquitous frataxin expression, FRDA pathology is tissue-specific, disproportionately affecting dorsal root ganglia sensory neurons, dentate nuclei of the cerebellum, corticospinal tracts and cardiomyocytes. The molecular basis for this selective vulnerability remains unresolved, suggesting that cell-type specific responses to frataxin deficiency shape disease susceptibility. This incomplete understanding is compounded by the lack of molecular biomarkers that capture FRDA biology beyond frataxin deficiency, thereby limiting therapeutic development and evaluation. Here, we integrated all available human bulk RNA-seq datasets in FRDA (23 datasets across 10 cell types), spanning disease-related (cardiomyocytes, sensory neurons) and relatively FRDA-spared cell types (fibroblasts, lymphoblastoid cells) under a unified analytical framework to identify transcriptional dysregulation underlying selective vulnerability and candidate biomarkers. Meta-analysis revealed recurrent transcriptional perturbations beyond *FXN*, involving long non-coding RNAs, translational control and cytoskeletal organisation. While shared transcriptional themes were observed, the specific biological programmes engaged were strongly cell-type dependent. The top candidate biomarkers, *MYH14, MEG9,* and *MEG8* showed preferential upregulation in disease-relevant cell types including sensory neurons and cardiomyocytes, supporting their potential relevance to selective vulnerability. Therapeutic responsiveness to these candidates were assessed across RNA-seq datasets from FRDA models exposed to diverse therapeutic strategies, including epigenetic modulation and *FXN*-targeting approaches, revealing that transcriptional alterations in FRDA are pharmacologically modifiable. To facilitate transparent exploration and reuse of these findings, we developed an interactive FRDA Transcriptomic Atlas, providing a community-accessible resource for investigating gene and pathway-level dysregulation across FRDA studies: https://marniemaddock.github.io/FRDATranscriptomicAtlas/. Together, these findings implicate cell type specific transcriptional programs as potential drivers of selective vulnerability and establish a framework for prioritising biomarkers in FRDA.

**Graphical Abstract:** 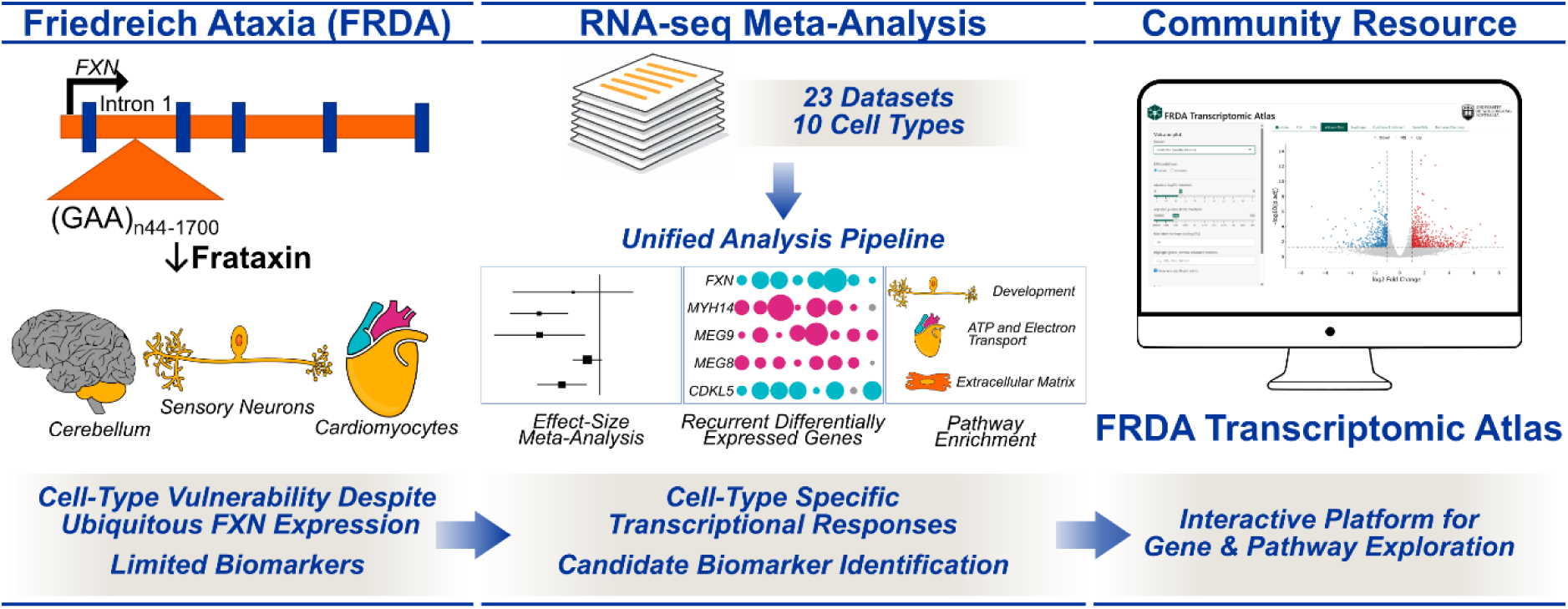

## Introduction

Friedreich ataxia (FRDA) is an autosomal recessive progressive neurodegenerative disorder primarily caused by a homozygous guanine-adenine-adenine (GAA) trinucleotide repeat expansion within the first intron of the frataxin (*FXN*) gene on chromosome 9 (1,2). The GAA expansion length varies between affected individuals, with reports of expansions up to 1700 repeats in FRDA, whilst unaffected individuals typically carry a GAA repeat of 5-34 (1,3). Age of onset and neurological severity is largely predicted by the size of the GAA expansion on the smaller allele (2). The GAA expansion in FRDA induces a transcriptionally repressive state, reducing expression of the protein product frataxin (4–7). Frataxin is essential for mitochondrial iron-sulfur cluster biogenesis, and its deficiency disrupts multiple mitochondrial pathways (8). Consequently, frataxin deficiency leads to oxidative stress (9–13), decreased activity of iron-sulfur cluster enzymes (14–16), decreased adenosine triphosphate (ATP) (17–21), loss of mitochondrial membrane potential (9,11,22) and iron accumulation (13,23). Although mitochondrial dysfunction is a central feature of FRDA, there is growing evidence that additional pathways, including lipid metabolism, the cytoskeleton and non-coding ribonucleic acid (RNA) networks are also dysregulated, highlighting the complexity of FRDA molecular pathology (24–28).

Frataxin is expressed in all cell types and is essential for life, as a complete loss is embryonically lethal in mice (29). Despite this ubiquitous expression, FRDA pathology is strikingly tissue specific (30). The dorsal columns of the spinal cord, corticospinal tract, dentate nuclei of the cerebellum, retinal ganglion cells, the dorsal root ganglia and peripheral sensory nerves, the heart, skeletal muscle and pancreas are disproportionately affected in FRDA (31–33). This clinically manifests as progressive sensory and proprioceptive loss, limb and gait ataxia, dysphagia, dysarthria, hearing and visual dysfunction, skeletal changes, muscle weakness and wasting, cardiomyopathy and metabolic dysfunction. The basis for this cell-type specific selective vulnerability, despite *FXN* deficiency in all cells, remains a central unanswered question in FRDA pathophysiology.

The first disease-modifying therapy Omaveloxolone was approved only in 2023 for affected individuals aged ≥ 16 years (34), 27 years after discovery of the genetic basis of FRDA (2). One reason therapeutic progress remains slow is the lack of reliable molecular biomarkers that capture FRDA biology beyond Frataxin deficiency. Current clinical outcome measures such as clinical rating scales (Modified Friedreich Ataxia Rating Scale (mFARS), Scale for the Assessment and Rating of Ataxia (SARA) or cardiac measures, have significant limitations including ceiling effects, require large sample sizes and require time to show meaningful change to demonstrate efficacy (35). Biochemical measures may offer faster outcomes that complement clinical outcomes; however, those commonly used in FRDA, including frataxin levels or oxidative damage markers such as urinary 8OH2ʹdG, capture specific aspects of the complicated FRDA disease biology.

*In vitro* assays face the similar limitations, as most therapeutic screens rely on frataxin expression or a single downstream pathway, most commonly mitochondrial function, as primary readouts. Such reductionist approaches may overlook therapies that improve broader cellular health without altering these mitochondrial readouts. Although multiple studies have reported FRDA cellular dysregulation across diverse pathways, these signals remain underutilised as potential biomarkers, prompting calls for a broader, multi-component biomarker “toolbox” for FRDA (35).

Ideally, molecular markers used *in vitro* should align with disease processes measurable *in vivo*, including both preclinical animal models and FRDA cohorts, while capturing multiple aspects of disease biology. Transcriptomic profiling is well suited to address this limitation, offering an unbiased and scalable approach to detect diverse FRDA-related dysfunction beyond frataxin and mitochondrial-centric readouts by capturing the whole transcriptome. Consistent with this, the 2018 FARA biomarker meeting highlighted omics approaches as promising strategies for identifying downstream disturbances overlooked by targeted assays (36).

Despite the growing number of transcriptomic and proteomic studies in FRDA, translation of differentially expressed genes (DEGs) and proteins into validated biomarkers has remained limited. The 2018 FARA biomarker report highlighted that many promising molecular biomarkers fail validation due to biological variability, model-specific effects, and lack of cross-study standardisation, emphasising the need for systematic, multi-model validation to distinguish core disease biology from context-dependent artefacts. This challenge is amplified by the marked tissue and cell-type specificity of FRDA pathology, whereby selective vulnerability of cardiomyocytes and sensory neurons contrasts with the relative sparing of other cell types such as fibroblasts. Although numerous bulk RNA sequencing (RNA-seq) datasets spanning diverse FRDA donors, tissues, and experimental models are now publicly available, these datasets have largely been analysed in isolation, limiting their ability to resolve shared versus cell-type-specific disease mechanisms.

Here, we investigated whether integration of publicly available human RNA-seq datasets across multiple FRDA cell types could reveal robust transcriptional signatures and candidate biomarkers. We performed a systematic review and meta-analysis of all currently available human bulk RNA-seq datasets in FRDA, comprising 23 datasets across 10 distinct cell types, including disease-relevant (cardiomyocytes, sensory neurons) and relatively FRDA-spared or later-affected cell-types (fibroblasts, lymphoblastoid cells, lower motor neurons). Using a unified analytical pipeline, we integrated these datasets to identify pan-FRDA molecular signatures and define cell-type specific drivers associated with selective vulnerability. This analysis revealed consistently dysregulated genes beyond *FXN*, highlighting recurrent transcriptional signatures involving long non-coding RNAs, cytoskeletal organisation, and translation, alongside pronounced cell-type specificity of these effects. Notably, the leading candidate biomarkers, *MYH14, MEG9,* and *MEG8* showed stronger dysregulation in disease-relevant induced pluripotent stem cell (iPSC)-derived sensory neurons and cardiomyocytes. To enable transparent exploration and reuse of these findings, we developed an interactive FRDA Transcriptomic Atlas, providing a community-accessible resource for exploring gene and pathway-level dysregulation across FRDA studies (37). We aim to update the Atlas as new bulk RNA-seq datasets from FRDA cohorts become available.

## Methods

### Systematic Review

This study followed the Preferred Reporting Items for Systematic Reviews and Meta-Analyses (PRISMA) guidelines to identify both records and datasets to be used as part of the systematic review and meta-analysis (Supplementary Table 1). An overview of the methods workflow is given in Figure 1.

**Figure 1.**
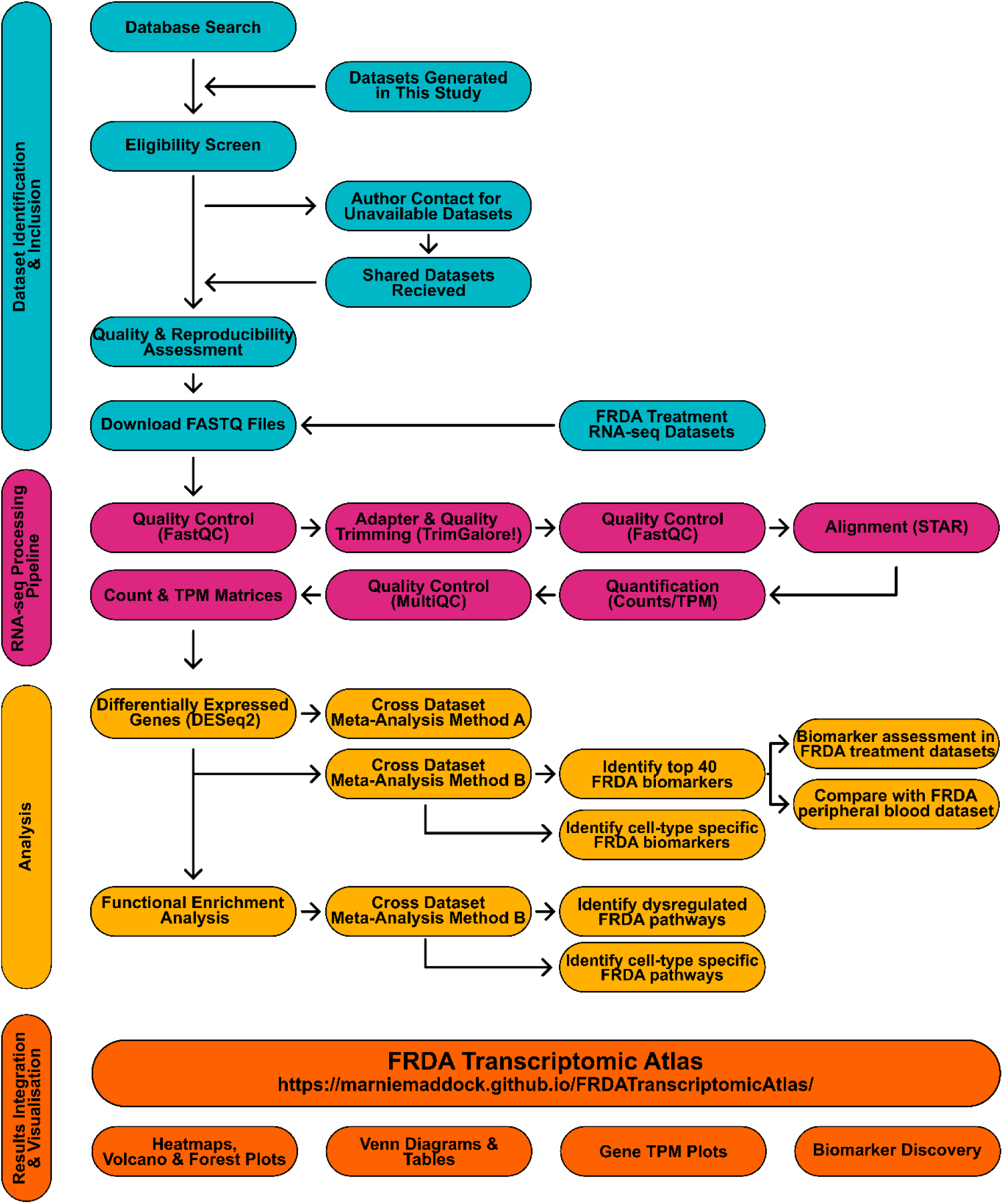
Overview of the systematic review and meta-analysis workflow, including dataset identification and inclusion, standardised RNA-sequencing processing, downstream analyses, and cross-dataset integration. All analyses derived from this meta-analysis, and additional visualisation options are available for exploration in the FRDA Transcriptomic Atlas app (37).

### Search Strategy

Records were searched from multiple sources, including Scopus, PubMed, the Gene Expression Omnibus (GEO), and BioRxiv, on September 5, 2025. This study focused on bulk RNA-seq data from FRDA samples to develop a search strategy. The terms used in Scopus, PubMed and BioRxiv were: (“Friedreich’s ataxia” OR “Friedreich ataxia” OR FRDA) AND (“RNA-Seq” OR RNAseq OR “RNA seq” OR “RNA sequencing” OR “RNA-sequencing” OR transcriptom*). The search strategy in GEO included “Friedreich’s ataxia”, limiting results to Homo sapiens and expression profiling by high-throughput sequencing. The exact queries used are described in Supplementary Table 2. No publication date restrictions were applied. All search terms were queried in English, which may have excluded relevant studies published in other languages. All identified studies were imported into Covidence for de-duplication, title and abstract screening, eligibility screening and data extraction.

### Dataset Selection

Records were first assessed for relevance at the title and abstract stage by one reviewer (MLM), with a second reviewer (AD) verifying the included and excluded records. Exclusion criteria at this stage were 1) studies not related to FRDA or 2) absence of RNA-seq data. Full-text articles were then screened using more stringent eligibility criteria. The inclusion criteria were as follows: 1) FRDA compared to a control group; 2) Human or Human-derived samples; 3) Bulk RNA-seq measurements; 4) raw sequencing data publicly available. The exclusion criteria included: 1) No FRDA case/control; 2) Non-human model; 3) FXN knockdown/overexpression; 4) No bulk RNA-seq data; 5) Raw sequencing data not publicly available; 6) Duplicate datasets (i.e. re-analysis of public sequencing data); 7) Other sequencing type (e.g. microarray, miRNA); 8) reverse transcriptase quantitative polymerase chain reaction (RT-qPCR) as the only transcriptomic approach; 9) Non-primary research (review, book chapter). The full-text review was performed by one author (MLM) and independently checked by a second reviewer (AD). Citation searching of the full-text articles was also performed to identify other relevant records. Unpublished datasets generated by our laboratory that met the inclusion/exclusion criteria were also included as part of this review (see Supplementary Methods 1). These included FRDA and isogenic corrected cell lines (38), differentiated into iPSC-derived neural crest cells, sensory neurons and lower motor neurons.

### Data Extraction

Nine studies met eligibility criteria and were included in the data extraction stage. For datasets that were not publicly available, corresponding authors were contacted to request access to the raw data for inclusion in this study, resulting in an additional two datasets. Data were extracted by one reviewer (MLM) and validated by another reviewer (AD). Although three studies focused on frataxin restoration or therapeutic testing, availability of raw FRDA and control data enabled the first direct cross-dataset comparison between these groups, yielding insights beyond the original study objectives (6,39,40). The data extraction stage included: 1) information collection on study design and characteristics, RNA-seq methods and reported results; 2) quality assessment comparable to a risk-of-bias evaluation; 3) obtain raw sequencing FASTQ files for standardised re-analysis.

To address the first component, we extracted data across the following categories: First Author (Year), Digit Object Identifier (DOI), Journal, Accession ID, Link to Dataset, Project Number, Sequencing type, Library Type, Sequencing Platform, RNA Extraction Method, RNA Integrity Number (RIN), Sample Source/Cell type), Sample Size, *FXN* expression, GAA repeat length, Number of Differentially Expressed Genes between FRDA and control, Up Genes (n), Down genes (n), Reported False Discovery Rate (FDR)/log_2_Fold-Change (FC) cutoffs, Key Pathway Enrichment Results.

To assess methodological quality and reproducibility, categories of information were extracted and rated on a three-point scale from 1 (low quality score), 2 (medium quality score) and 3 (high quality score). The criteria used to score each category are given in Table 1. These criteria were informed by established best practices (41) or by shared expertise and agreement within the research team.

**Table 1:**
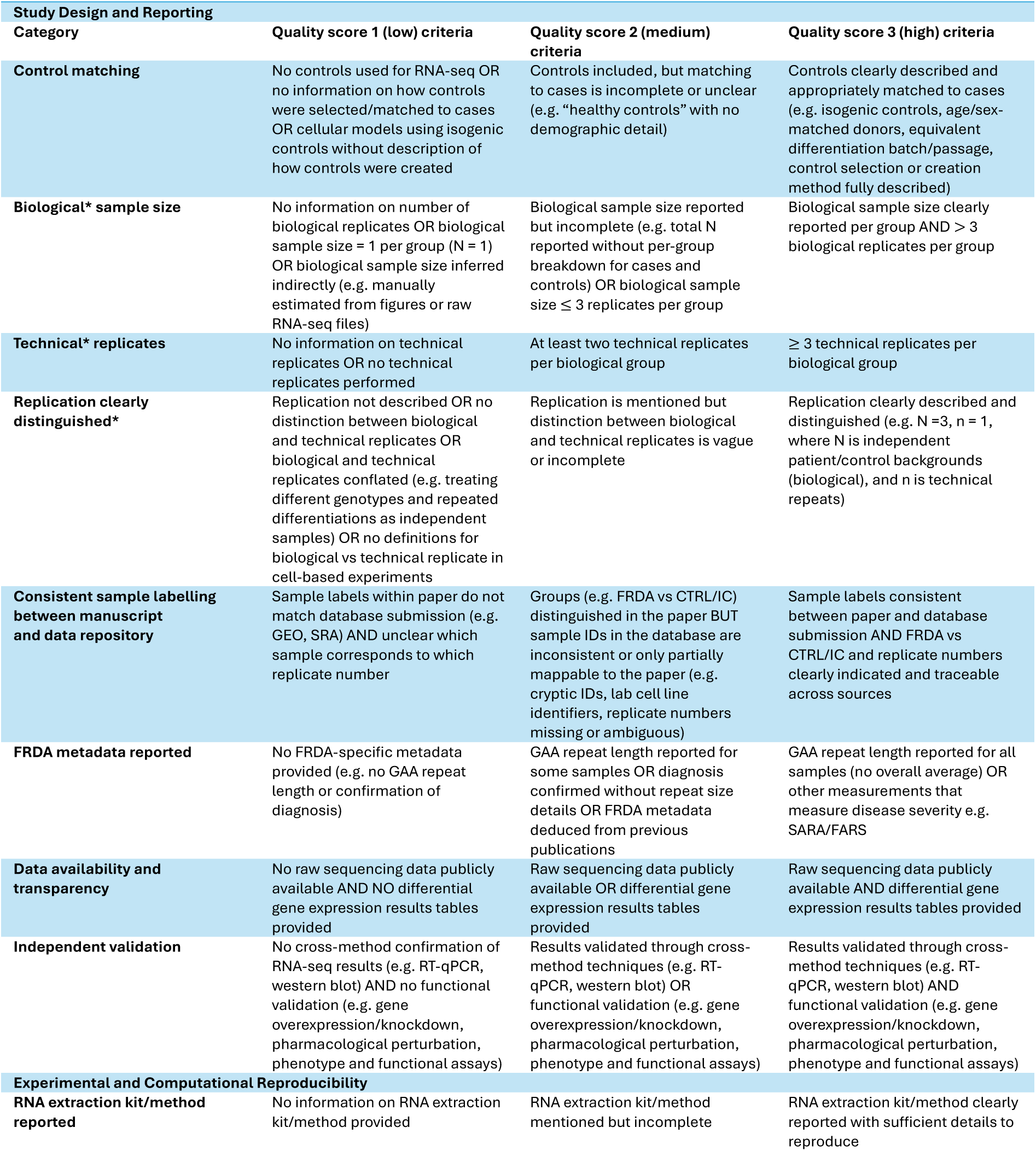
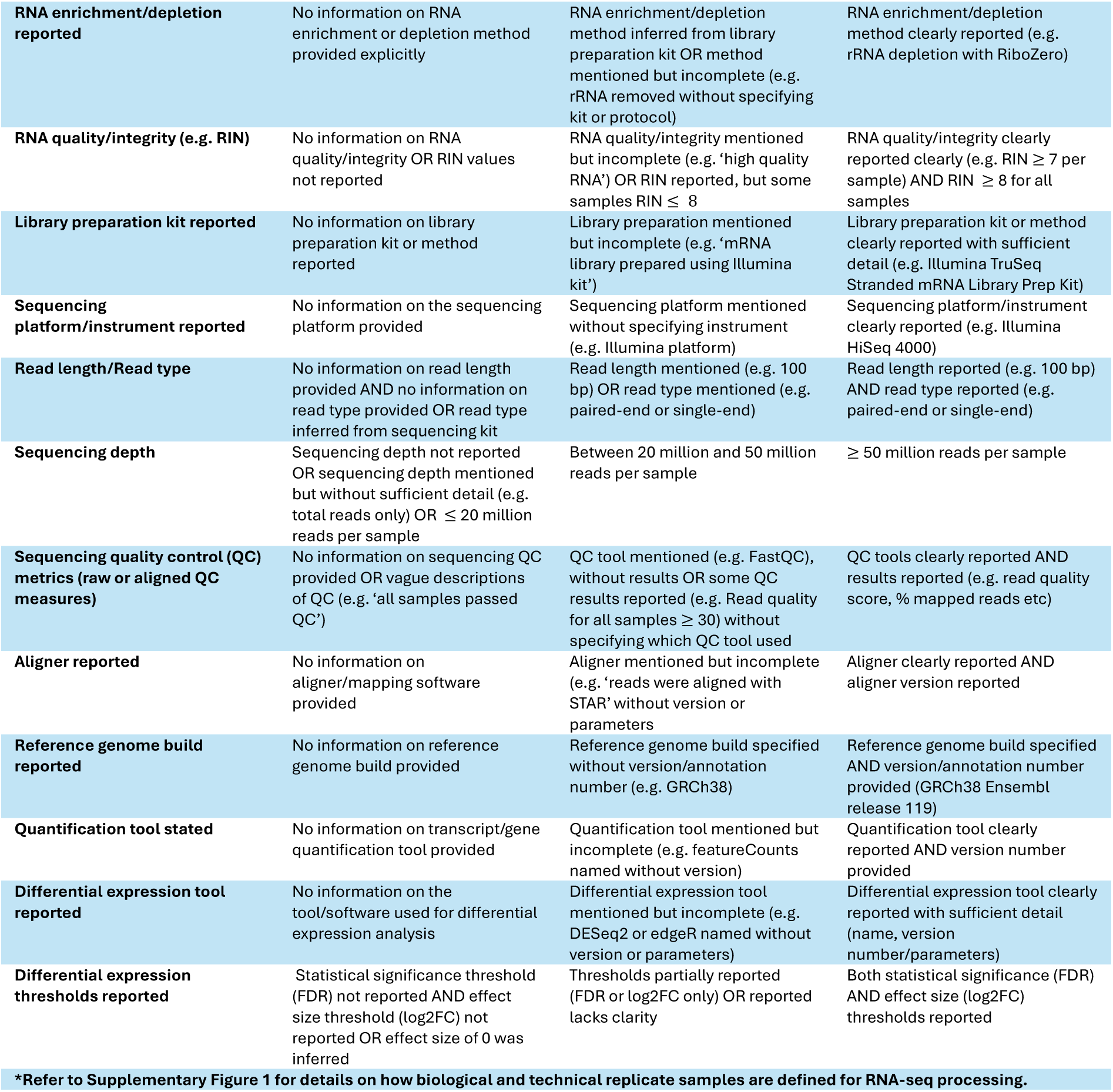
Methodological quality and reproducibility criteria definitions.

The categories to assess methodology quality and reproducibility were subdivided into 1) study design and reporting, and 2) experimental and computational reproducibility (Table 1). Each study had an overall mean score computed from all the categories. In addition, mean scores were calculated for each category to identify specific aspects of reporting that were weaker and may benefit from improvement in future FRDA RNA-seq studies. Values are given as mean (standard deviation) (SD).

### Meta-Analysis - RNA Sequencing Processing Pipeline

All datasets included in this meta-analysis with available raw FASTQ files from bulk RNA-seq were processed using the same computational workflow. This also applied to samples generated in our laboratory and are published within this study (Supplementary Methods 1). Raw FASTQ files were processed using the nf-core RNA-seq pipeline (version 3.18.0) (42). Initial read quality of FASTQ files was assessed using FastQC (v0.12.1). Adapter and quality trimming was performed using TrimGalore! (v0.6.10; cutadapt v4.9) with default parameters which automatically detect and remove adapter sequences, trim low-quality bases (Phred < 20) and remove short reads (< 20 bp). Trimmed reads were aligned to the human reference genome (GRCh38, Ensembl release 113) using STAR (v2.8.11b). Gene and transcript level quantification was performed using Salmon (v1.10.3) to generate raw count matrices and transcripts per million (TPM) normalised expression values. Alignment and further quality control metrics were summarised and checked with MultiQC.

### Per-dataset Differential Expression

Differentially expressed genes and transcripts between FRDA and control samples were analysed per dataset with DESeq2 v1.48.1 (43) using raw count matrices. Genes/transcripts with < 10 total reads across all samples were excluded before normalisation and model fitting. For each dataset, variance-stabilised transformation counts were assessed using principal component analysis (PCA), sample-to-sample Euclidean distance heatmaps, and dispersion-mean plots. Outlier samples that consistently failed to cluster with their biological group in both PCA and distance analyses were removed. This included one FRDA sample within the Erwin dataset (Supplementary Figure 2). When datasets showed evidence of internal technical batches (i.e., samples clustering together independent of biological group), a batch term was incorporated into the DESeq2 design formula (design = ∼ batch + group). For datasets with multiple donor pairs (e.g., *Lees*), differential expression was computed separately for each donor comparison between individuals with FRDA and controls (e.g., FA1 vs FA1ic, FA2 vs FA2ic, FA3 vs FA3ic).

*FXN* expression was assessed across different cell-types within the same study using log_2_ transformed TPM values (log_2_(TPM+1)). All analyses were performed within individual studies stratified by the cell line (isogenic corrected or FRDA) and analysed separately. Differences in *FXN* expression across cell types were assessed using one-way ANOVA. Model assumptions were assessed using the Shapiro-Wilk test for residual normality and Levene’s test for homogeneity of variance (all *p* > 0.05). Tukey HSD test was used for pairwise comparisons. Data are presented as mean log_2_(TPM + 1) (SD).

### Cross-Dataset Differential Expression Meta-Analysis – Meta-analysis Method A

To identify genes showing consistent average effect size changes across individuals with FRDA versus control contrasts, we performed a pooled meta-analysis of differential expression estimates. Meta-analyses of differential expression were conducted by collating all DESeq2-derived log_2_FC estimates and standard errors across FRDA versus control contrasts. For each gene, a random effects meta-analysis was fitted using restricted maximum likelihood in the metafor package (rma.uni) in R, giving a pooled effect size, standard error, z-statistic and I^2^. P values were FDR adjusted across genes. Genes were retained for visualisation purposes if they were present in at least 19 datasets, had an absolute pooled log_2_FC > 0.5, FDR < 0.05, and directional agreement exceeding 65%. Directional agreement was quantified as the proportion of contributing datasets in which the gene’s log_2_FC was positive or negative.

### Differential Expression Overlap Analysis Across Datasets – Meta-Analysis Method B

As meta-analysis method A prioritises effect size magnitude and direction at the gene level, it may be less suited to capturing transcriptional changes that are consistently significant across datasets but variable in magnitude between experimental contexts or cell types. To complement the pooled meta-analysis, we also performed an overlap analysis of DEGs to quantify the recurrence of dysregulated genes across FRDA datasets and different cell-types. For each dataset, genes with FDR ≤ 0.05 were separated into upregulated (log_2_FC > 0) and downregulated (log_2_FC < 0) sets. These were then counted to determine the number of datasets in which each gene was significantly up or downregulated. Genes detected as differentially expressed in more than eight datasets (approximately one third of datasets) were retained for visualisation to prioritise reproducible signals across datasets.

### Candidate Biomarker Panel Identification

Based on the most consistently up or downregulated genes in FRDA, we defined a candidate transcriptomic biomarker panel comprising the top 40 ranked genes. A full ranked list of all genes are given in Supplementary File 2. Genes were prioritised using a hierarchical ranking method that favoured genes with (i) the highest number of studies showing significant up or downregulation overall, (ii) the fewest conflicting studies in terms of significant up versus downregulation, iii) stronger directional agreement as quantified by the direction score, and (iv) larger absolute median log_2_ FC as a final tie-breaker. The direction score ranges from -1 (consistently downregulated) to +1 (consistently upregulated) with values near zero indicating weak or inconsistent directionality. Extended methodological details describing this strategy are given in Supplementary Methods 2. The top 40 genes were then evaluated in an independent peripheral blood microarray dataset from 418 individuals with FRDA and 93 unaffected controls to assess their potential relevance as blood-based biomarkers (44). To identify cell-type specific biomarkers and pathways, analyses were additionally performed separately for major cell types, including fibroblasts (Napierala, Vilema-Enriquez, Wang), lymphoblastoid cells (Erwin, Chutake), cardiomyocytes (Lees, Li), sensory neurons (Lai - PNS, Maddock - SN), CNS/LMNs (Lai - CNS, Maddock - LMN) and all neurons (Lai (CNS/PNS), Maddock (SN/LMN), Mishra). The full cell-type specific biomarker panels are given in Supplementary File 2.

### Functional Enrichment Analysis - GSEA

Functional enrichment analysis was performed for each dataset using gene set enrichment analysis (GSEA) on ranked gene lists with the gseGO function from the clusterProfiler package in R. Enrichment was assessed independently for the three gene ontology (GO) categories biological process, molecular function and cellular component. Only gene sets containing 10 to 500 genes were included. Pathways with an FDR < 0.05 were considered significantly different between FRDA and control samples. For each ontology, the top 15 pathways were selected for visualisation based on the number of datasets in which the pathway showed significant enrichment in FRDA versus controls. A ranked list of GSEA results was also generated for major cell type groups based on the number of datasets showing significant enrichment, directional consistency, and median normalised enrichment score (NES). The full cell-type stratified results are given in Supplementary File 3. Redundant GO terms were collapsed using semantic similarity-based reduction (rrvgo package; similarity threshold 0.7), retaining representative parent pathways for visualisation of the top 10 positively and negatively enriched terms. Complete enrichment results for all individual datasets are available through the FRDA Transcriptomic Atlas (37).

### Treatment Datasets

To evaluate the behaviour of consistently dysregulated genes under therapeutic intervention, we analysed publicly available FRDA treatment RNA-seq datasets. These analyses provide proof-of-concept pharmacodynamic readouts, assessing whether consistently dysregulated genes respond to therapeutic intervention, rather than demonstrating treatment efficacy. These datasets included FRDA primary lymphocytes treated with 10 mM nicotinamide (Chan: GSE42960) (45), lymphoblastoid cell lines from FRDA individual and a healthy sibling control treated with a synthetic transcription elongation factor (syn-TEF1) designed to allow transcription along repressed GAA expansions (Erwin: GSE99403) (39), FRDA fibroblasts treated with an inhibitor of histone methyltransferase SUV20-H1 (A-196, 1 µM, 5 µM, 10 µM) (Vilema-Enriquez: GSE145115) (46), FRDA fibroblasts treated with antisense oligonucleotides targeting *FXN* (S30, S10, S10+2, S10_L6) (Wang: GSE205526) (40), skeletal muscle biopsies from individuals with FRDA treated with recombinant human Erythropoietin (rhuEPO) (Indelicato: GSE226646) (47), and FRDA iPSC-derived neurons with the histone deacetylase inhibitor 109 (HDACi 109) (Lai: PRJNA495860) (48). Each study included matched untreated or vehicle-treated controls (e.g., untreated control, vehicle control, siNTC), enabling a clear FRDA treatment versus FRDA control contrast within the same donor backgrounds. Raw sequencing files were processed with the same pipelines described above. Differential expression was computed for each treatment condition relative to its matched FRDA control condition to assess treatment-associated transcriptional changes.

### FRDA Transcriptomic Atlas

The FRDA Transcriptomic Atlas was developed as an interactive application to enable exploration and cross-study comparison of FRDA transcriptomic data analysed in this meta-analysis. The application provides access to full differential expression and functional enrichment results, as well as visualisation tools including PCA plots, volcano plots, tables, heatmaps and Venn diagrams. Datasets can be directly compared within the application using Venn diagrams and forest plots. Individual genes of interest can also be queried and visualise transcript abundance as TPM values. Installation instructions are given here: (37).

## Results – Systematic Review

### Dataset Selection

Database searching identified 144 records, with an additional 44 records coming from citation searching or grey literature (Figure 2). After removal of duplicates, 121 unique records were screened by title and abstract. Of these, 53 were excluded for not relating to FRDA or not including RNA-seq data. The remaining 68 records were assessed for eligibility by reading the full-text articles. A total of 59 records were excluded; 17 for not having a human model (e.g. *yeast* or *Drosophila*), 13 records with duplicate datasets (i.e. re-analysis of publicly available datasets, 7 records performing RNA-seq, but were not using bulk methods (e.g. microarray or miRNA), 5 studies that did not perform RNA-seq, 3 non-primary research articles, 2 studies using *FXN* knockdown or overexpression, 2 studies had non-FRDA populations, and 2 were excluded for not including any FRDA cases with controls. There were 8 records that would have passed all inclusion and exclusion criteria apart from their raw FASTQ files not being publicly available. Therefore, we contacted authors of these 8 studies and requested their data for inclusion in this study. An additional 2 datasets were obtained from authors which were included in this study. Overall, this yielded 11 studies that were included in this review and meta-analysis.

**Figure 2:**
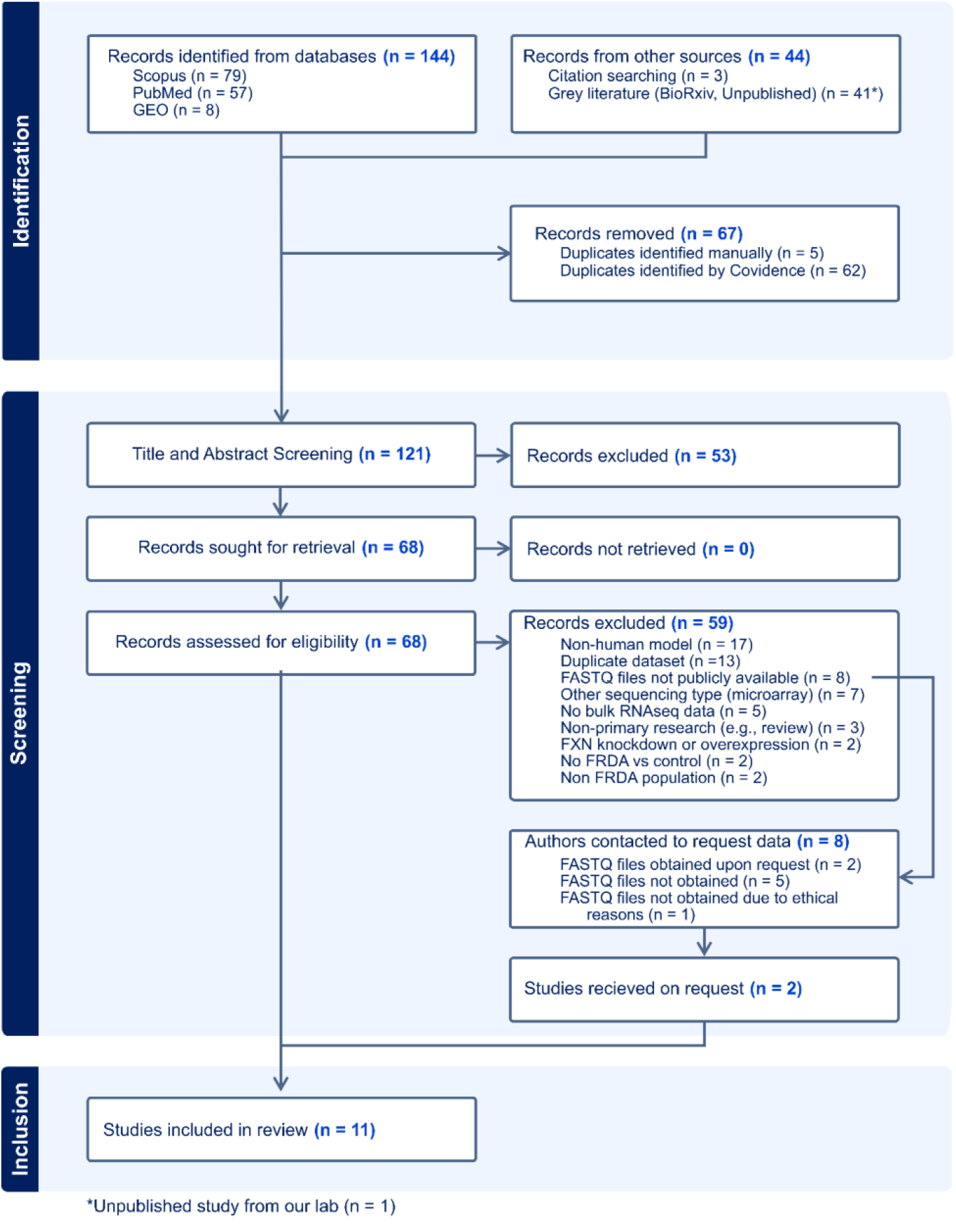
PRISMA flow chart summarising the identification, screening, exclusion, and inclusion of studies containing human FRDA transcriptomic datasets in this systematic review and meta-analysis.

### Study Characteristics

In total, 11 studies were eligible for data extraction, covering publications from 2014 to 2026. Among these, one was a preprint available on BioRxiv (49), and another was unpublished datasets from our laboratory that met eligibility criteria. From these 11 studies, there were 23 distinct RNA-seq datasets available for analysis. Study characteristics are summarised in Table 2. A metadata file mapping FASTQ files to samples is given in Supplementary File 4. These 23 datasets collectively represented 94 FRDA and 99 control samples, with individual study sizes ranging from 1 to 18 FRDA donor backgrounds with corresponding controls. Sample sources comprised of patient-derived lymphoblastoid cells (6,39), skeletal muscle (47), fibroblasts (40,46,50) and several iPSC FRDA or control-derived populations including iPSCs, peripheral nervous system sensory (PNS) neurons and central nervous system (CNS) neurons (48), sensory neurons (SNs), lower motor neurons (LMNs), neural crest cells, neurons (51) and cardiomyocytes (49,52). This covers a broad range of cell types that are typically affected in FRDA pathology (e.g. cardiomyocytes, sensory neurons) as well as cell types less directly associated with disease (e.g. fibroblasts, LMNs). Notably, no post-mortem FRDA brain or spinal cord samples were identified, with all data derived from primary or iPSC-based models. All FRDA samples were homozygous for GAA expansion, which is the most common FRDA pathogenic variant.

**Table 2:**
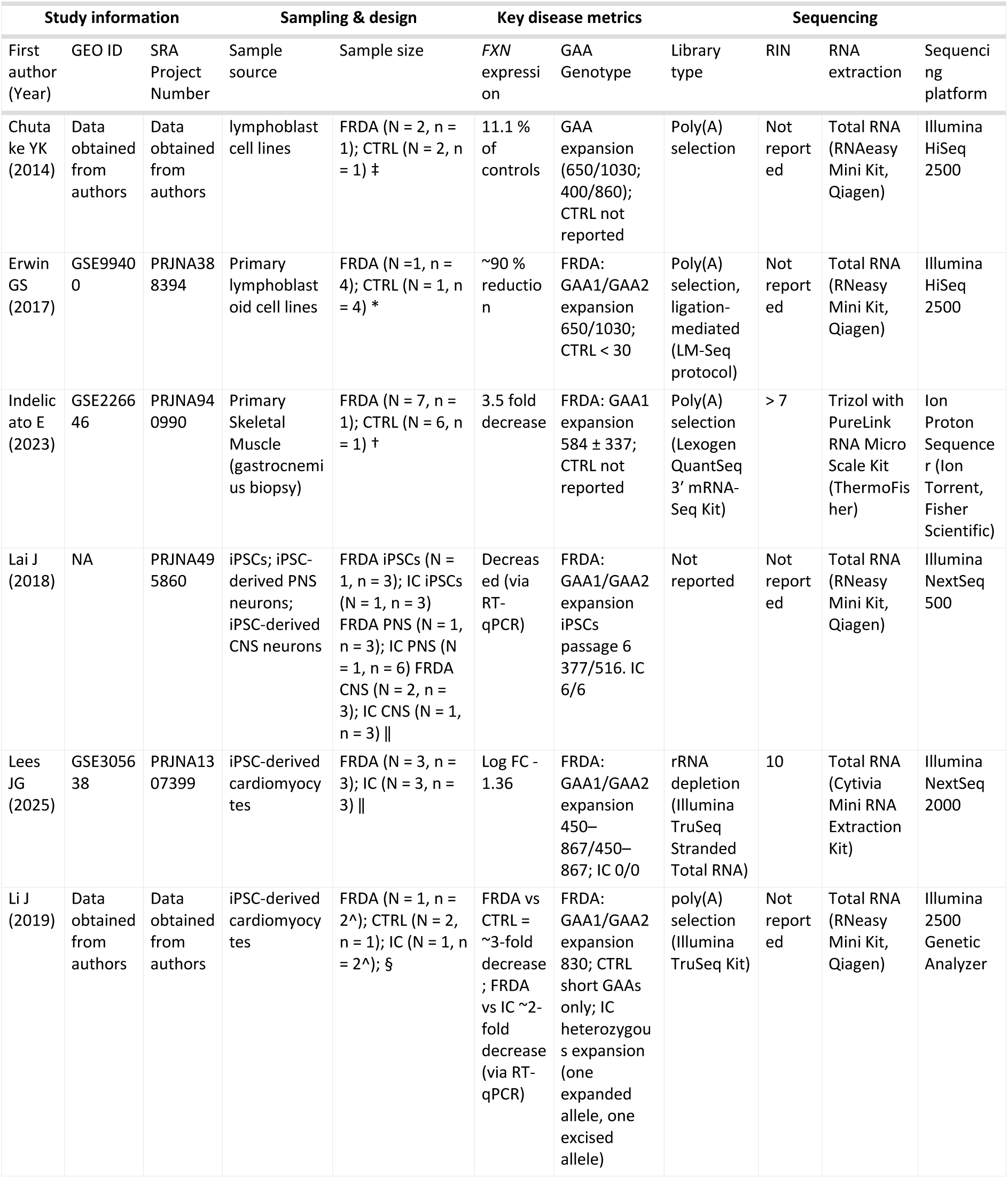
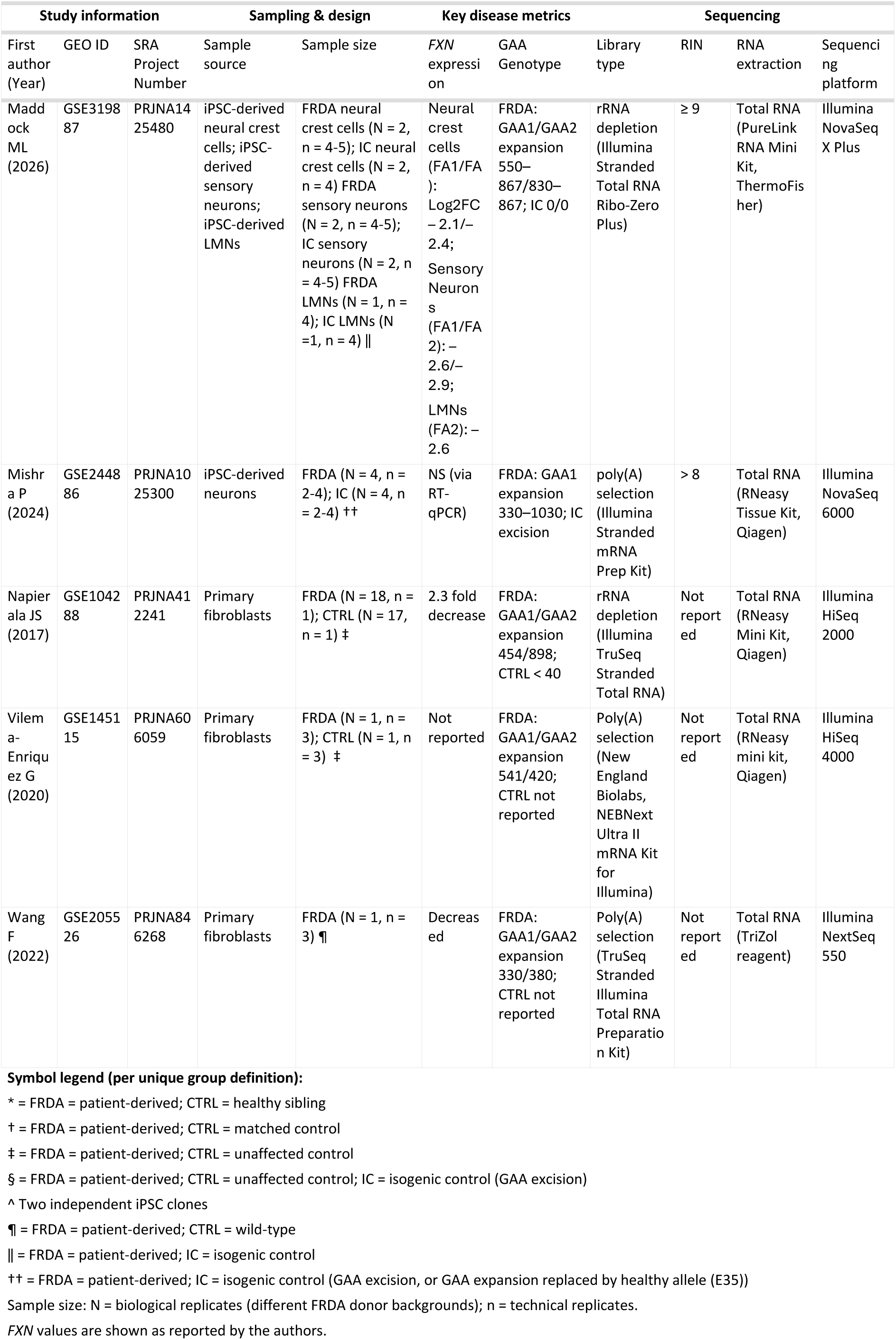
Included RNA-seq studies: Study Characteristics.

All included studies performed bulk RNA-seq. RNA extraction methods varied across the included studies, although all used commercially available kits. Library preparation was based on poly(A) tail selection in 7 studies (6,39,40,46,47,51,52), ribosomal RNA (rRNA) depletion in three studies (42,44, this study), while one study did not specify the method used (48). Sequencing was carried out predominantly on Illumina platforms (HiSeq, NextSeq, NovaSeq), although one study used the Ion Proton Sequencer (Ion Torrent) from Fisher Scientific (47). This heterogeneity in RNA extraction and sequencing methodology is a potential source of technical variability and is a consideration for downstream analyses. RNA quality, read length, read type and sequencing depth were inconsistently documented (Table 2, Figure 4). These gaps highlight challenges in comparing datasets and underscore the importance of re-analysis under a standardised pipeline.

### Quality and Reproducibility Assessment

We assessed the quality and reproducibility of included studies on numerous categories from a scale of 1 (low quality) to 3 (high quality). The specific definitions for each category’s quality criteria are given in Table 1 and quality scores for each study and category are shown in Figure 3. The overall average score for all studies across all categories was 2.24 (0.48). The highest scoring category across all studies was ‘RNA Extraction Kit’, with a perfect score of 3.0, followed by ‘sequencing platform/instrument reported’ and ‘FRDA metadata reported’ with average scores of 2.91 (0.30). The lowest scoring categories were ‘RNA quality/integrity (e.g. RIN)’ and ‘Sequencing depth’ with average scores of 1.64 (0.92) and 1.73 (0.65) respectively. When aggregating categories into 1) study design and reporting and 2) experimental and computational reproducibility, the average scores across all studies are 2.24 (0.35) and 2.33 (0.36) respectively. This highlights that RNA-seq methods were often better documented than the overall study design and reporting. Altogether, this identifies key areas across FRDA RNA-seq studies that can be improved upon in future studies to increase reproducibility and transparency standards.

**Figure 3:**
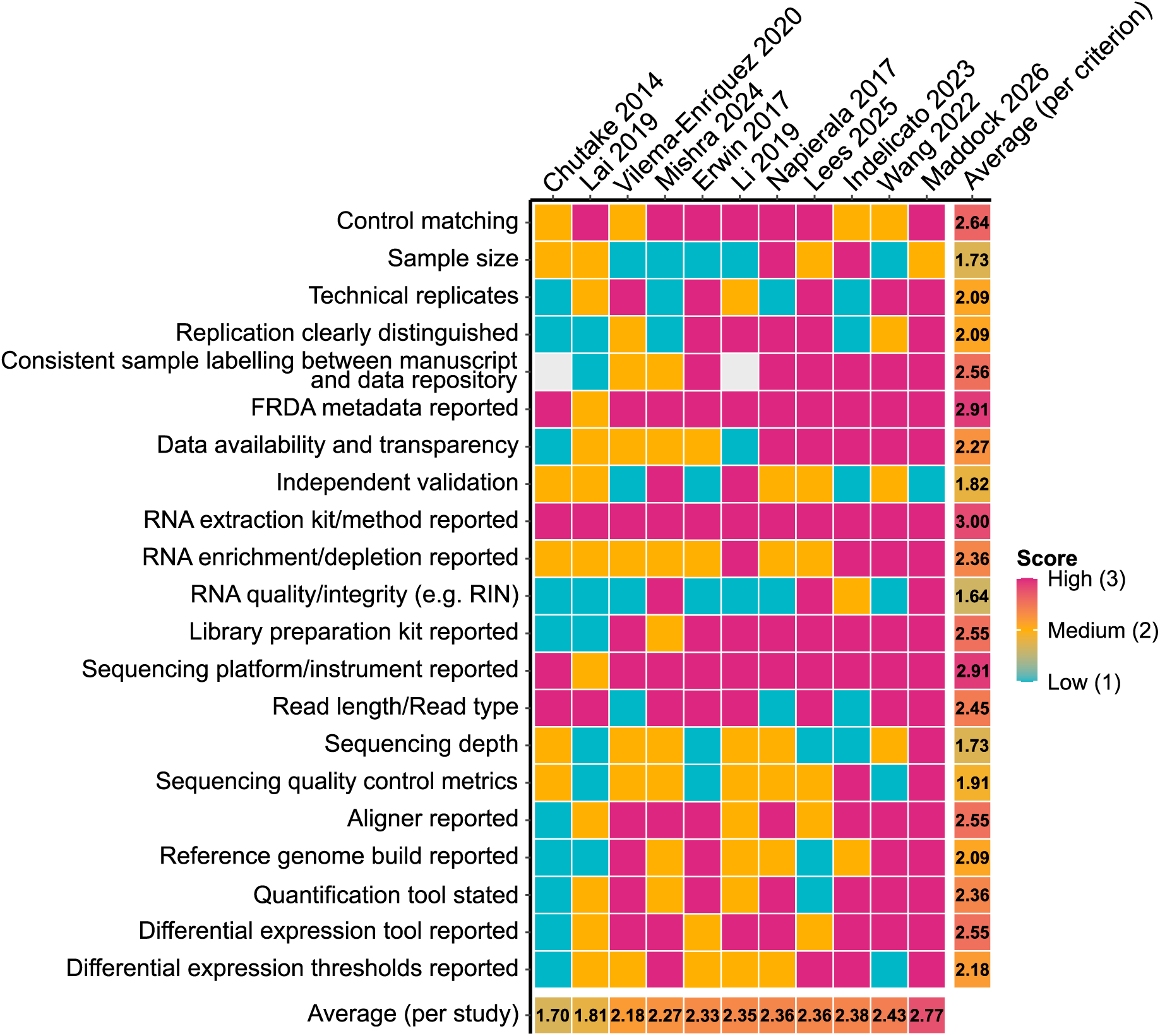
Study quality assessment scores across transcriptomic datasets. Each cell represents the score assigned to a given study for a specific criterion: low = 1 (blue), medium = 2 (yellow), high = 3 (pink). Scores were independently assigned based on predefined criteria by two reviewers and resolved by consensus.

With advances in RNA-seq technologies and growing awareness of reporting standards, we examined whether study quality improved over time. Spearman’s correlations showed no significant correlation between publication year and overall study quality scores (*ρ* = 0.59, *p* = 0.06) or ‘study design and reporting’ (*ρ* = 0.40, *p* = 0.23). However, there was a significant positive correlation between publication year and ‘experimental and computational reproducibility’ (*ρ* = 0.67, *p* = 0.02), suggesting improvements in methodological reporting and transparency over time.

## Results – Meta-Analysis

### Sequencing Metrics

All studies were processed through the same standardised RNA-seq analysis pipeline. The quality of the raw sequencing data, as well as other sequencing metrics such as median read length, depth of sequencing and unique mapping rate, were assessed to ensure comparability across datasets. All studies had excellent raw base-calling accuracy after trimming, with sample averages ranging between 92.8% to 100% of bases exceeding a Phred score above 30, corresponding to a ≥ 99.9% base-calling accuracy (Figure 4A). The median read length varied across studies, ranging from 50 ± 0 bases (52) to ∼ 166 ± 19 bases (47) (Figure 4B). Whilst longer reads generally improve mapping specificity, unique mapping percentages remained high in all datasets, with rates of 82.6 ± 8.5 % (51) to 93.1 ± 0.5 % (49) (Figure 4C). Sequencing depth showed the greatest variation across studies, with the lowest average depth observed in the Erwin (39) (4.7 ± 1.7 million reads) and Indelicato study (47) (5.2 ± 1.3 million reads), and the highest in the dataset generated for this study by our group (124.2 ± 33.5 million reads) (Figure 4D). All of these sequencing measures show that in spite of technical differences, every study generated high-quality sequencing data that is suitable for downstream analysis.

**Figure 4.**
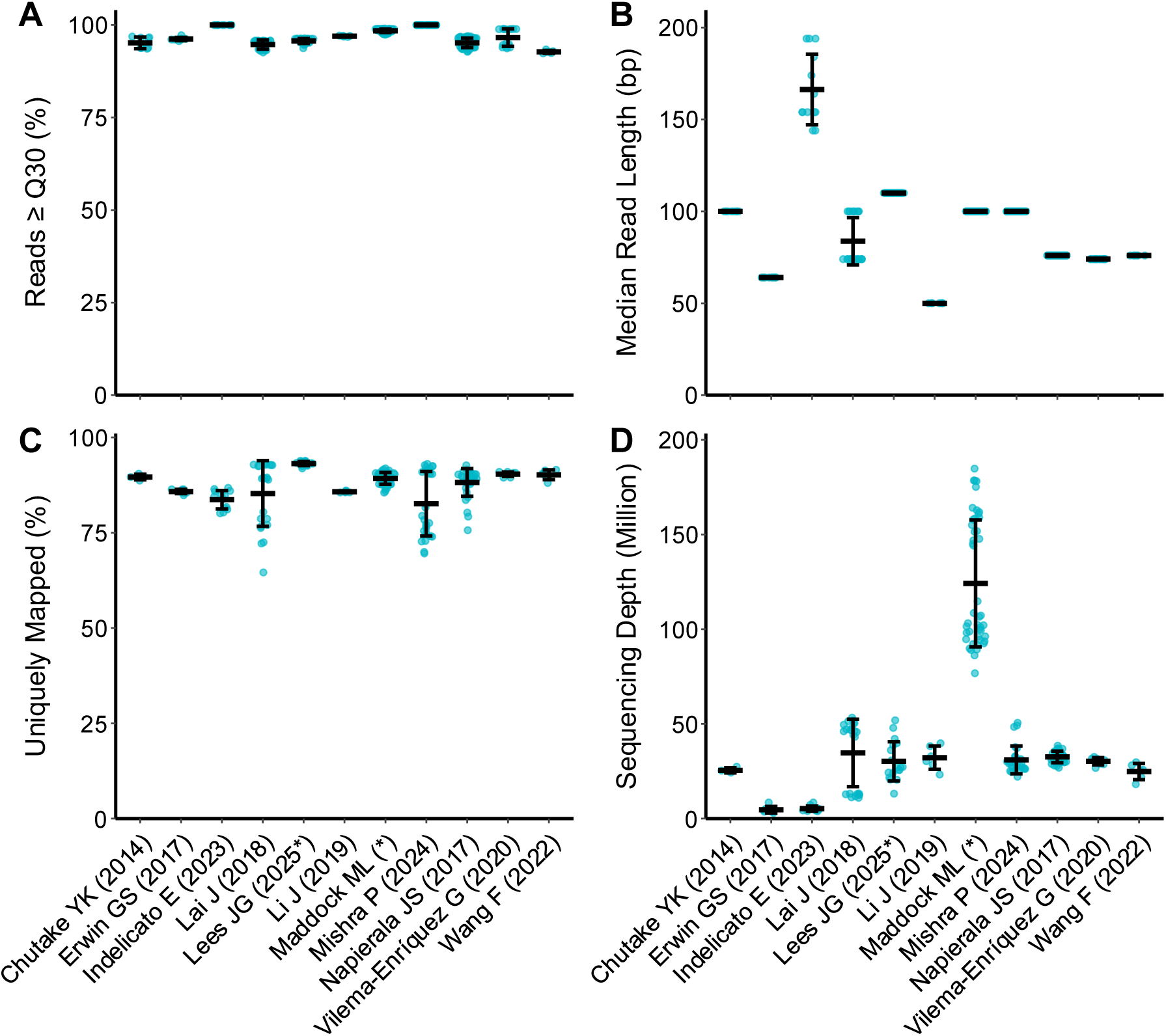
Sequencing quality and coverage metrics across FRDA RNA-seq datasets. A) Percentage of bases with a Phred quality score ≥ 30. B) Median read-length (base pairs) per dataset. C) Percentage of uniquely mapped reads. D) Sequencing Depth (millions of reads per sample). Each point represents individual samples from that dataset; black bars indicate study-level mean ± standard deviation. All datasets met quality thresholds appropriate for downstream comparative analyses. *Studies on BioRxiv at time of analysis or new datasets generated in this study.

### Variation in the Number of Differentially Expressed Genes across FRDA Cell-types and Donors

To assess transcriptional dysregulation across FRDA cell types, we quantified the number of DEGs in each study using the same analysis pipelines. In instances where studies included multiple patient-derived lines with repeated differentiations from the same FRDA donor, each comparison was quantified separately to preserve donor-specific backgrounds. Table 3 summarises the number of up and downregulated genes (FDR < 0.05), as well as those meeting a log_2_ fold change of ≤ -1 or ≥ 1. The total number of DEGs varied between studies and cell types (between 171 and 10812 total DEGs). There was high variability within the same cell type across studies. For example, primary FRDA fibroblasts from the Napierala, Vilema-Enriquez and Wang studies had 4409, 3208 and 941 DEGs, respectively, compared to controls. The number of DEGS also differed within cell types in the same study. For example, in iPSC-derived cardiomyocytes from Lees et al., one patient line had only 224 DEGs, while another had 6705 compared to the isogenic corrected controls. This was also evident in iPSC-derived neural crest cells (FA1: 1275 DEGs, FA2: 4345 DEGs) and sensory neurons (FA1: 3256 DEGs, FA2: 8612), highlighting substantial patient-specific differences. These findings underline the complexity of FRDA transcriptomics, with both experimental conditions and patient background contributing to DEG variability.

**Table 3:**
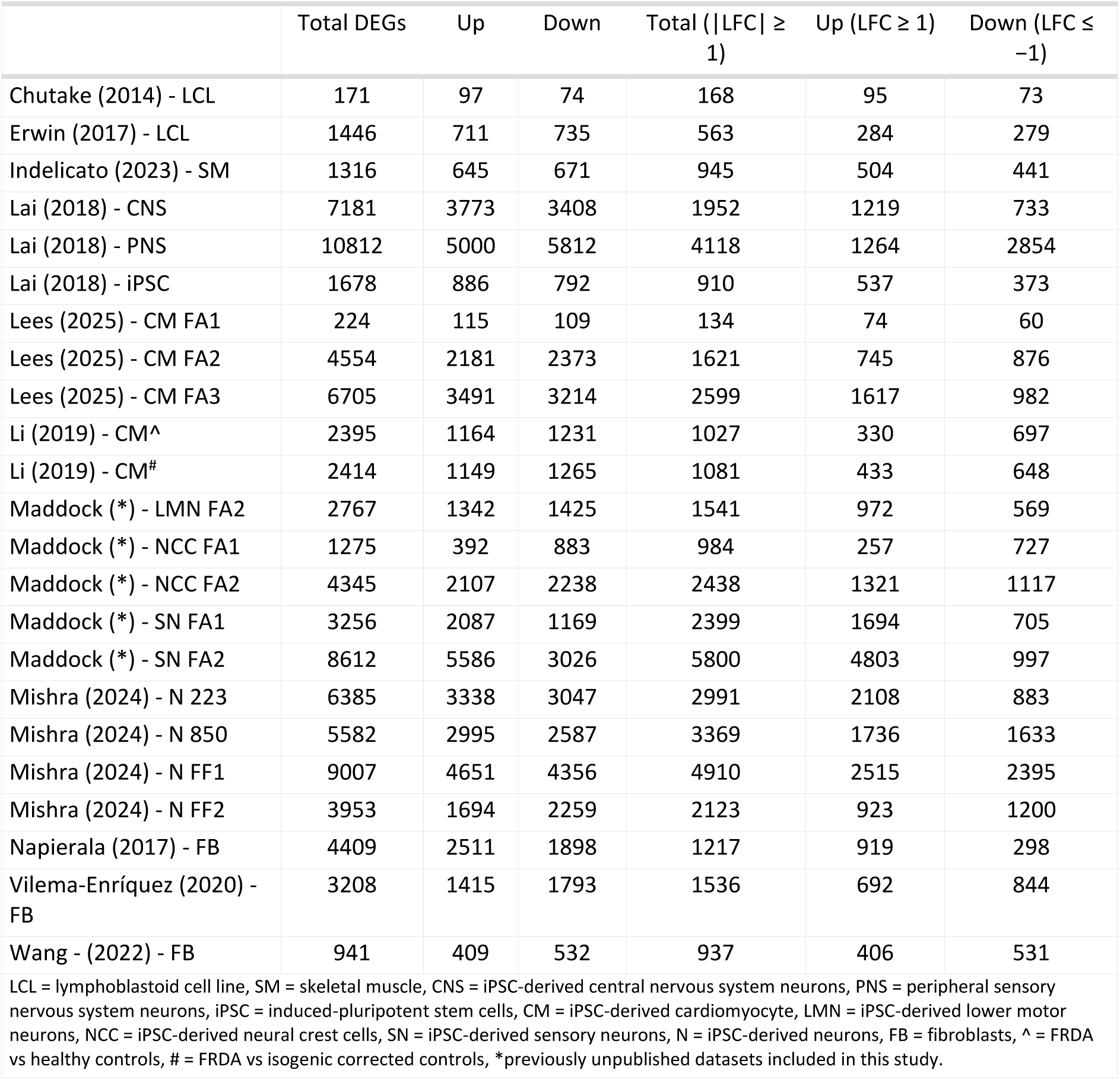
Differentially Expressed Genes (n) (FDR ≤ 0.05)

A unique aspect of collating these datasets is that we can compare transcriptomic changes across multiple cell types derived from the same donor. The iPSC cell lines FA1 and FA2 with corresponding isogenic controls FA1ic and FA2ic were differentiated into cardiomyocytes, neural crest cells and sensory neurons, while the FA2/FA2ic line additionally generated lower motor neurons. DEG numbers for FA1 varied by cell type, with cardiomyocytes showing the least (224 DEGs), followed by neural crest cells (1275 DEGs) and then sensory neurons (3256 DEGs), highlighting that the degree of transcriptional disruption in FRDA is shaped by cellular context. Interestingly, the FA2 line showed the fewest DEGs in lower motor neurons (2767 DEGs), a cell type whilst impacted by FRDA, are usually considered secondary in FRDA pathology. This was followed by neural crest cells (4345 DEGs), then cardiomyocytes (4554 DEGs) and finally sensory neurons (8612 DEGs). This pattern may indicate relative resilience of motor neurons to *FXN* deficiency compared with sensory neurons or cardiomyocytes, which are initially more apparent in the pathology of FRDA. Although sequencing depth and library complexity may partially explain variation in DEG numbers, the divergent patterns in DEGs in different cell types between FA1 and FA2 lines suggest technical factors alone are unlikely to fully account for these findings. These differences may reflect patient-specific patterns of cell type vulnerabilities that contribute to phenotypic heterogeneity. Further, Lai et al., performed RNA-seq on iPSCs, CNS neurons and PNS sensory neurons from the same cell lines (48). Again, a cell type that is prevalent and impacted early in FRDA pathology, peripheral sensory neurons, had the largest number of DEGs (10812) in FRDA compared to isogenic controls, followed by CNS neurons (7181), then iPSCs (1678). Inclusion of iPSCs suggests that transcriptional alterations become more pronounced upon differentiation into affected cell types, hinting at a possible developmental aspect to FRDA pathogenesis. Together, these highlight the importance of considering both cellular context and patient-specific factors when interpreting transcriptional dysregulation in FRDA.

### *FXN* Expression

Given that reduced *FXN* expression is the defining molecular defect in FRDA, it serves as an internal benchmark for confirming dataset validity. A forest plot of per-dataset and pooled meta-analytic log_2_ fold change showed strong downregulation of *FXN* in nearly all datasets (Figure 5A, C). Overall, the random effects model confirmed a significant reduction in *FXN* (log_2_FC = -1.51, 95% CI: -1.92, -1.10; I^2^ = 95.5%, Figure 5C). Notably, the Mishra study, which generated neurons from four different FRDA patient iPSCs and compared them to isogenic corrected cells, did not show consistent downregulation of *FXN*. One comparison (line 223) even suggested increased *FXN* expression (log_2_FC = 1.11, 95% CI: 0.72, 1.50, Figure 5C), while the other three comparisons showed no evidence of differential expression, with effect sizes close to zero. This likely reflects variability relating to technical factors, gene-editing efficiency or intrinsic cell line behaviour. However, these FRDA cell lines displayed a significant decrease in Frataxin expression at the protein level (51), indicating that transcript levels may not fully reflect functional protein deficiency. Similarly, fibroblasts from Wang et al., also showed no significant differences in *FXN* between FRDA and controls, however had high variability (log_2_FC = -0.74, 95% CI: -1.54, 0.05, Figure 5C). All other studies showed a negative log_2_ fold change in *FXN* expression between FRDA and controls. Although *FXN* expression from Mishra et al. 2024 and Wang et al., 2022 was not significantly decreased, FRDA-related cellular phenotypes may arise from the GAA expansion itself, including epigenetic and transcriptional alterations extending beyond *FXN* expression. These datasets were therefore included in downstream analyses.

**Figure 5:**
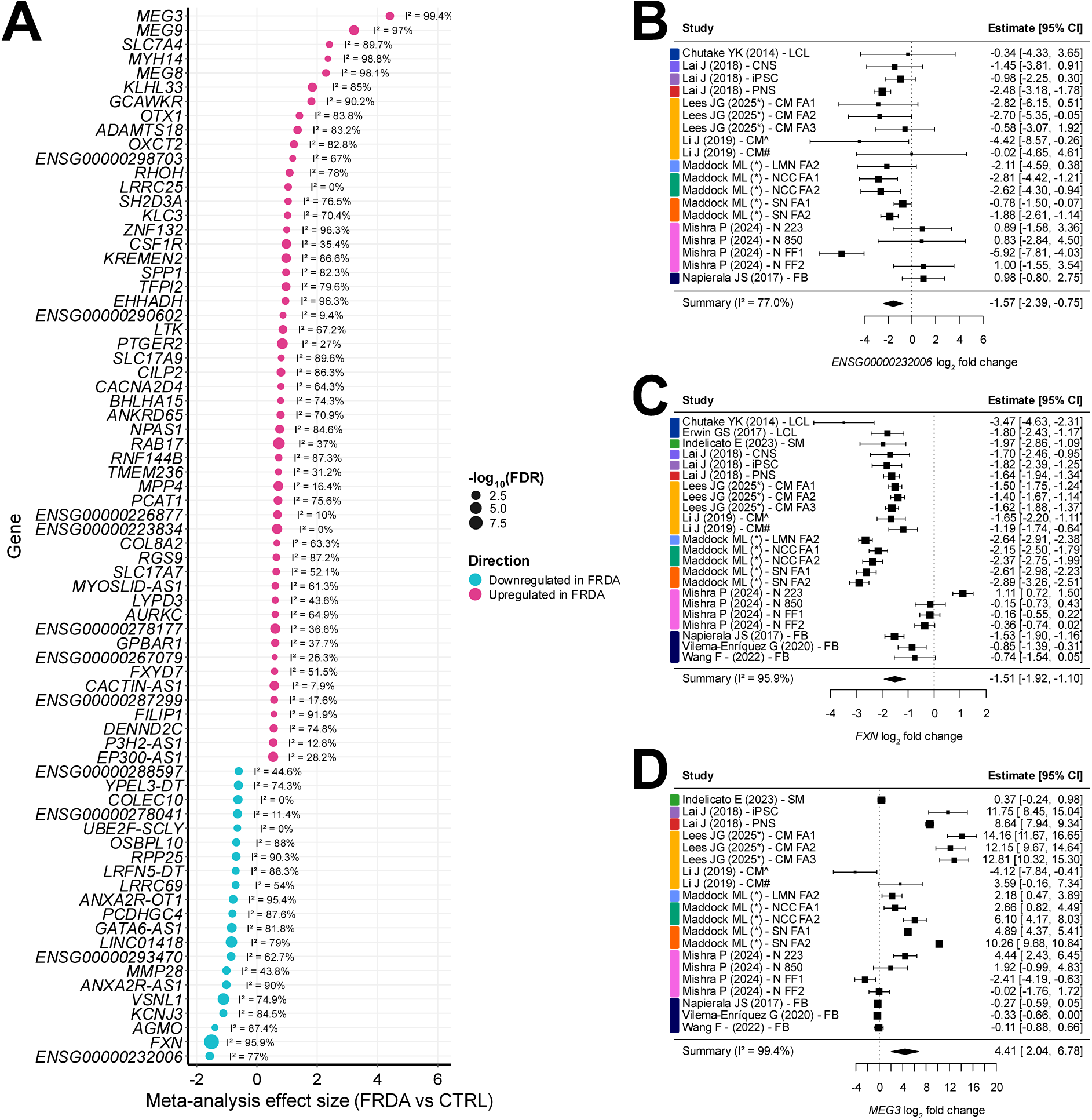
Effect-size meta-analysis identifies core FRDA-associated genes with substantial heterogeneity (Meta-Analysis Method A). A) Random effects meta-analysis of log2 fold-change across all FRDA vs control datasets. Each point represents the pooled effect size (β), scaled by -log10(FDR) and coloured by direction (FRDA vs control; pink = upregulated, blue = downregulated). I^2^ values denote between-dataset heterogeneity for that gene across all datasets. Displayed genes met predefined inclusion thresholds (gene measured in ≥ 19 datasets, |β| > 0.5, FDR < 0.05m, > 65 % directional concordance). B-D) Forest plots for the topmost overall downregulated gene *ENSG00000232006* (B), the causal gene in FRDA *FXN* (C), and the topmost overall upregulated gene *MEG3* (D). Abbreviations: FRDA = Friedreich ataxia, LCL = lymphoblastoid cell line, SM = skeletal muscle, CNS = iPSC-derived central nervous system neurons, PNS = peripheral sensory nervous system neurons, iPSC = induced-pluripotent stem cells, CM = iPSC-derived cardiomyocyte, LMN = iPSC-derived lower motor neurons, NCC = iPSC-derived neural crest cells, SN = iPSC-derived sensory neurons, N = iPSC-derived neurons, FB = fibroblasts, ^ = FRDA vs healthy controls, # = FRDA vs isogenic corrected controls, *Studies on BioRxiv at time of analysis or new datasets generated in this study.

To assess whether *FXN* expression varied across cell types, we compared log₂(TPM + 1) values across different cell types derived from the same cell lines within the Lai and Maddock datasets. In the Lai datasets, *FXN* expression differed significantly across cell types in both isogenic corrected (F(2,9) = 355.8, *p* < 0.0001) and FRDA cells (F(2,6) = 68.4, *p* < 0.0001), with CNS neurons showing consistently lower expression (IC: 0.93(0.05); FRDA: 0.29(0.19)) than PNS sensory neurons (IC: 4.04(0.16), *p* < 0.0001; FRDA: 2.30(0.37), *p* < 0.0001) and iPSCs (IC: 4.72(0.31), *p* < 0.0001; FRDA: 3.03(0.31), *p* < 0.0001). There was a significant reduction in *FXN* expression between isogenic corrected PNS neurons and iPSCs (*p* < 0.01), but not between FRDA PNS neurons and iPSCs (*p* = 0.054). In contrast, when comparing multiple cell types derived from the same genetic background in the Maddock (FA2) dataset, *FXN* expression did not differ significantly across cell types (neural crest cells, sensory neurons, LMNs) in the isogenic corrected cell line (F(2,21) = 0.49, *p* = 0.62), and only a modest effect was observed in FRDA cells across the three cell-types (F(2,9) = 5.00, *p* = 0.03), driven by a single significant reduction in *FXN* between LMNs (1.99(0.30)) and neural crest cells (2.70(0.30), *p* = 0.034). Collectively, these findings indicate that *FXN* expression varies across cell types within individual studies and cell lines; however, the inability to directly compare TPM values across datasets limits cross-study interpretation of these patterns.

### Effect-size Based Meta-analysis (Method A) Identifies Core FRDA Genes with Substantial Cross-study Heterogeneity

To identify which genes were consistently dysregulated across FRDA versus CTRL contrasts, a meta-analysis was performed on the per dataset log_2_FC estimates and standard errors per gene. There were 74 genes that had a combined meta-analysis log2FC (FRDA vs CTRL) estimate (β) > 0.5, FDR < 0.05, measured in ≥ 19 datasets and > 65 % directional concordance (i.e. > 65 % of datasets were significantly up or downregulated for that gene (Figure 5A). The uncharacterised long non-coding RNA *ENSG00000232006* showed the largest magnitude of downregulation in the meta-analysis (β = -1.57, SE = 0.41, FDR = 8.5 x 10^-3^, I^2^ = 77 %; Figure 5A, 5B), followed by *FXN* (β = -1.51, SE = 0.21, FDR = 1.8 x 10^-10^, I^2^ = 95.9 %; Figure 5A, 5C), consistent with its deficiency in FRDA. The long non-coding RNA *MEG3* showed the largest overall magnitude of upregulation in FRDA vs control (β = 4.41, SE = 1.21, FDR = 1.1 x 10^-4^, I^2^ = 99.4 %; Figure 5A, 5D). The remaining top genes encompassed several functional categories, including long non-coding RNAs (*ANXA2R-AS1*, *ANXA2R-OT1*, *GATA6-AS1*, *LINC01418*, *LRFN5-DT*, *ENSG00000278041*, *YPEL3-DT*, *ENSG00000288597*, *EP300-AS1*, *ENSG00000223834*, *MYOSLID-AS1*, *PCAT1, MEG9, MEG8*), transcription factors (*NPAS1, OTX1, BHLHA15, ZNF132*), neuronal signaling and synaptic function (*PCDHGC4, NPAS1, SLC17A7, SLC17A9, KCNJ3, RGS9, FXYD7, VSNL1, KLC3, CACNA2D4, MPP4*), lipid transport or metabolism (*OSBPL10, CILP2, OXCT2, AGMO, EHHADH*), RNA processing (*RPP25*), and vesicle trafficking (*RAB17*, *DENND2C*). However, between-dataset heterogeneity was substantial for most genes (I² > 50 % in 52/74 genes), indicating that the magnitude of differential expression varies markedly across datasets. For this reason, we applied a complementary recurrence-based meta-analysis (Method B) to assess the consistency of differential expression across FRDA datasets.

### Recurrent Upregulated Genes Across FRDA Datasets

To characterise transcriptional changes that recur across FRDA datasets independent of effect-size magnitude, we ranked genes according to the number of FRDA versus control comparisons in which they were significantly dysregulated using meta-analysis method B. This recurrence-based approach highlights transcriptional alterations consistently observed across studies and enables assessment of cell-type contributions to the between-dataset heterogeneity identified in meta-analysis method A (Figure 6). The top upregulated gene across all datasets (13/23) was *MYH14,* which encodes the non-muscle myosin II-C (NMIIC) motor protein (Figure 6). Interestingly, *MYH14* was significantly increased in FRDA cells coming from affected FRDA cell types including PNS sensory neurons, sensory neurons, cardiomyocytes and in unresolved cell-types including iPSCs, neural crest cells (FDR < 0.05), but not fibroblasts, CNS neurons, lower motor neurons, skeletal muscle or lymphoblastoid cells. Significant upregulation of *MYH14* was also observed in FRDA iPSC-derived neurons that did not display significant decreased *FXN* expression (Mishra datasets). Recurrent upregulation of *MYH14* in disease-relevant cell types supports its association with FRDA-related cellular processes and as a candidate biomarker of disease involvement in FRDA.

**Figure 6:**
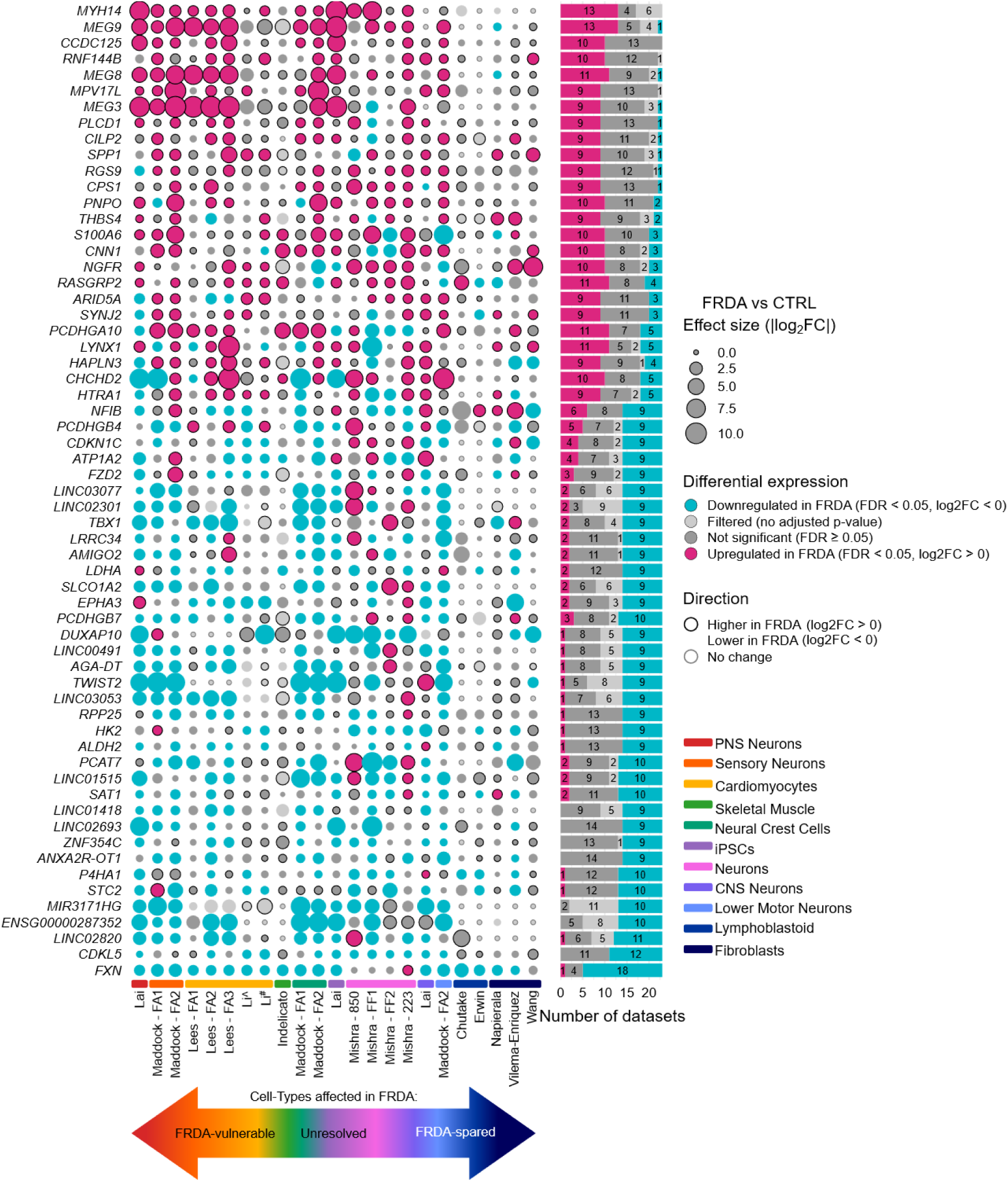
Cross-dataset recurrence analysis of differentially expressed genes in FRDA (meta-analysis method B). Bubble plot showing gene-level differential expression across each FRDA vs control dataset. Genes differentially expressed in more than 8 studies are plotted. The size of the point corresponds to absolute log_2_ fold-change (FRDA vs control). For visualisation purposes, values were capped at the 99th percentile to limit the influence of extreme outliers on point scaling. Colours indicate significance (FDR < 0.05) and direction (pink: log2FC > 0, blue: log2FC < 0). For non-significant genes (FDR ≥ 0.05), points are coloured dark grey, and have a black outline if the gene was expressed higher in FRDA compared to controls, or no outline if expressed lower in FRDA compared to controls. Genes with no adjusted p-value due to independent filtering in DESeq2 are coloured light grey. The bar plot summarises the number of datasets supporting upregulation, downregulation, or no significant change for each gene. ^ FRDA vs healthy control, ^#^ FRDA vs isogenic corrected control.

The next most consistently significantly upregulated gene across all datasets was the long non-coding RNA *MEG9* which was upregulated in 13/23 FRDA versus control comparisons (Figure 6). *MEG9* belongs to a locus on chromosome 14 called the *DLK1-DIO3* locus. Two additional long non-coding RNAs from this locus, *MEG8* and *MEG3*, were also upregulated in 11/23 and 9/23 FRDA versus control comparisons, respectively. *MEG9* was upregulated in PNS neurons, cardiomyocytes, and sensory neurons which are known cell types affected in FRDA pathology, but not in CNS neurons, fibroblasts, or lymphoblastoid cells. *MEG9* was also increased in skeletal muscle, another pathological cell type - in FRDA vs control cells (log_2_FC = 4.12), however showed variability between donors, resulting in no adjusted *p* value following multiple corrections testing. Interestingly, cell types of unknown significance in FRDA pathology, such as iPSCs and neural crest cells also displayed this increased *MEG9* expression in FRDA compared to controls, consistent with early-stage or differentiation-linked regulation at the *DLK1-DIO3* locus. *MEG8* (upregulated in iPSCs, PNS neurons, cardiomyocytes, lower motor neurons, sensory neurons) and *MEG3* (upregulated in iPSCs, PNS neurons, cardiomyocytes, and sensory neurons) showed similar enrichment in disease-relevant cell types. Altogether this implicates *MEG9, MEG8* and *MEG3* as potential biomarkers in cell types affected in FRDA pathology and highlights coordinated transcriptional dysregulation involving the *DLK1-DIO3* locus, although whether these changes represent causal drivers or downstream consequences of disease remains unclear.

### Recurrent Downregulated Genes Across FRDA Datasets

Alongside *FXN*, the next most consistently downregulated differentially expressed gene across FRDA versus control comparisons was *CDKL5*, downregulated in 12/23 datasets (Figure 6). *CDLK5* encodes a serine/threonine protein kinase, and *CDKL5* deficiency is a cause of the neurological condition *CDKL5* deficiency disorder. Interestingly, there was no uniform cell-type specific patterning of *CDKL5*. The most consistent reductions were observed in FRDA PNS neurons, with variable findings between donor backgrounds of iPSC-derived sensory neurons, cardiomyocytes and neural crest cells, and no differences in FRDA versus controls in lymphoblastoid cell lines and fibroblasts. The next most recurrent significantly downregulated DEGs across FRDA versus control comparisons were predominantly non-coding RNAs. The long intergenic non-coding RNAs (lincRNAs) *LINC02820, LINC02693, LINC01418, LINC01515, LINC03053, LINC00491, LINC02301, LINC03077* and the long non-coding RNAs (lncRNAs) *PCAT7, MIR3171HG*, *ENSG00000287352, ANXA2R-OT1, AGA-DT* were each reduced in FRDA in 9 to 11 of the 23 datasets. These genes showed no strong cell-type specific pattern, suggesting a broader disruption of regulatory RNA networks in FRDA.

### Prioritisation and Peripheral Validation of a 40-gene Candidate Biomarker Panel

To integrate the most consistently upregulated and downregulated genes identified across datasets, a combined ranked transcriptomic biomarker panel comprising the top 40 genes was generated (Table 4). To evaluate the clinical utility of prioritised transcriptomic biomarkers, we then compared our panel with a large peripheral blood microarray study of 418 individuals with FRDA and 93 unaffected controls. A total of 6 of 40 genes were significantly dysregulated in both peripheral blood and the meta-analysed datasets, however, directionality was not always consistent. Notably, *RNF144B*, an E3 ubiquitin ligase with diverse roles in proteasomal degradation and apoptosis regulation emerged as a promising biomarker, showing consistent dysregulation in peripheral blood and complete directional agreement across all datasets (directionality score = 1.00). *RNF144B* may represent a clinically tractable biomarker that bridges peripheral readouts with molecular pathology in disease-relevant tissues. A full ranked list of all genes across all datasets and cell-type stratified candidate biomarkers are given in Supplementary File 2.

**Table 4:** Prioritised FRDA candidate biomarkers based on recurrence of differential expression across transcriptomic datasets and median log2 fold change, with corresponding differential expression in an independent FRDA peripheral blood dataset shown for comparison.

| Rank | Gene | Class | Direction in FRDA vs Ctrl (majority) | Direction Score | Median log<sub>2</sub>FC | Peripheral Blood log<sub>2</sub>FC - FRDA vs control (sig status) |
| --- | --- | --- | --- | --- | --- | --- |
| 1 | *FXN* | protein-coding | down | -0.89 | -1.64 | NA |
| 2 | *MYH14* | protein-coding | up | 1.00 | 1.31 | 0.00904 (ns) |
| 3 | *MEG9* | lncRNA | up | 0.86 | 3.43 | -0.00462 (ns) |
| 4 | *CDKL5* | protein-coding | down | -1.00 | -0.40 | NA |
| 5 | *MEG8* | lncRNA | up | 0.83 | 1.51 | -0.0239 (sig) |
| 6 | *LINC02820* | lncRNA | down | -0.83 | -1.17 | NA |
| 7 | *RASGRP2* | protein-coding | up | 0.47 | 0.56 | -0.0602 (sig) |
| 8 | *PCDHGA10* | protein-coding | up | 0.38 | 1.00 | -0.0011 (ns) |
| 9 | *LYNX1* | protein-coding | up | 0.38 | 0.85 | NA |
| 10 | *ENSG00000287352* | lncRNA | down | -1.00 | -4.21 | NA |
| 11 | *MIR3171HG* | lncRNA | down | -1.00 | -3.19 | NA |
| 12 | *RNF144B* | protein-coding | up | 1.00 | 0.74 | 0.0873 (sig) |
| 13 | *CCDC125* | protein-coding | up | 1.00 | 0.43 | 0.0767 (ns) |
| 14 | *STC2* | protein-coding | down | -0.82 | -0.56 | 0.00139 (ns) |
| 15 | *P4HA1* | protein-coding | down | -0.82 | -0.32 | 0.0119 (ns) |
| 16 | *PCAT7* | lncRNA | down | -0.67 | -0.89 | NA |
| 17 | *LINC01515* | lncRNA | down | -0.67 | -0.77 | NA |
| 18 | *SAT1* | protein-coding | down | -0.67 | -0.28 | 0.0665 (sig) |
| 19 | *PNPO* | protein-coding | up | 0.67 | 0.24 | -0.0192 (ns) |
| 20 | *CNN1* | protein-coding | up | 0.54 | 1.43 | NA |
| 21 | *S100A6* | protein-coding | up | 0.54 | 0.94 | 0.0521 (sig) |
| 22 | *NGFR* | protein-coding | up | 0.54 | 0.77 | -0.000043 (ns) |
| 23 | *PCDHGB7* | protein-coding | down | -0.54 | -0.54 | 0.0082 (ns) |
| 24 | *CHCHD2* | protein-coding | up | 0.33 | 0.12 | -0.0674 (sig) |
| 25 | *LINC01418* | lncRNA | down | -1.00 | -1.09 | NA |
| 26 | *MPV17L* | protein-coding | up | 1.00 | 0.79 | -0.000663 (ns) |
| 27 | *ANXA2R-OT1* | lncRNA | down | -1.00 | -0.44 | NA |
| 28 | *LINC02693* | lncRNA | down | -1.00 | -0.31 | NA |
| 29 | *ZNF354C* | protein-coding | down | -1.00 | -0.19 | -0.00528 (ns) |
| 30 | *MEG3* | lncRNA | up | 0.80 | 3.12 | 0.00952 (ns) |
| 31 | *TWIST2* | protein-coding | down | -0.80 | -1.22 | 0.00527 (ns) |
| 32 | *AGA-DT* | lncRNA | down | -0.80 | -1.03 | NA |
| 33 | *SPP1* | protein-coding | up | 0.80 | 1.03 | 0.0198 (ns) |
| 34 | *RPP25* | protein-coding | down | -0.80 | -0.99 | 0.00342 (ns) |
| 35 | *LINC00491* | lncRNA | down | -0.80 | -0.95 | NA |
| 36 | *DUXAP10* | lncRNA | down | -0.80 | -0.93 | NA |
| 37 | *LINC03053* | lncRNA | down | -0.80 | -0.73 | NA |
| 38 | *RGS9* | protein-coding | up | 0.80 | 0.70 | NA |
| 39 | *CILP2* | protein-coding | up | 0.80 | 0.67 | NA |
| 40 | *CPS1* | protein-coding | up | 0.80 | 0.50 | 0.00714 (ns) |

Direction Score ranges from -1 to +1, where values close to -1 or +1 indicate strong directional consistency across studies, and values near 0 indicate mixed or inconsistent direction. Median log<sub>2</sub>FC between FRDA and controls were calculated across all datasets including non-significant datasets.
Direction Score = (number of upregulated studies - number of downregulated studies) / total significant studies.
NA = not available (gene not measured), ns = non-significantly different between FRDA and controls, sig = significantly different between FRDA and controls.

### Gene Set Enrichment Analysis Identifies Recurrent Translation and Cytoskeletal Signatures in FRDA

Gene set enrichment analysis was performed independently for each dataset to identify biological pathways, cellular compartments or molecular functions enriched in FRDA cells relative to controls. We then assessed whether common pathways were enriched across all datasets (Figure 7). The most recurrently enriched term was ribosome (GO:0005840), which showed significant enrichment in 20/23 datasets, including 8 with positive (NES > 0) and 12 with negative (NES < 0) scores. A cluster of recurrently enriched terms that were related to translation and the ribosome was observed across all datasets. Significant enrichment of terms related to protein synthesis machinery, including ribosomal subunit (GO:0044391), large ribosomal subunit (GO:0015934), cytosolic ribosome (GO:0022626), cytosolic large ribosomal subunit (GO:0022625), structural constituent of ribosome (GO:0003735), ribonucleoprotein complex biogenesis (GO:0022613) and cytoplasmic translation (GO:0002181) was observed across both FRDA affected and unaffected cell types. Contractile muscle fiber (GO:0043292) also showed significant enrichment in 19/23 datasets, including 13 with positive (NES > 0) and 6 with negative (NES < 0) scores. This GO term includes multiple genes that converge on cytoskeletal functions involved in cell shape, adhesion, contraction, intracellular transport and mechanotransduction (53), pointing towards dysregulation of cytoskeletal integrity in FRDA. This is further supported by the enrichment of additional cytoskeleton-related terms, including focal adhesion (GO:0005925), actin binding (GO:003779), heparin binding (GO:0008201), microtubule binding (GO:0008017), tubulin binding (GO:0015631), myofibril (GO:0030016) and cell-substrate junction (GO:0030055). Further, enrichment of the synaptic signaling pathways; regulation of trans-synaptic signaling (GO:0099177) and modulation of chemical synaptic transmission (GO:0050804) suggest that dysregulation of cytoskeletal components may extend to synaptic compartments, potentially impacting neuronal communication. Other signaling pathways may also be implicated in FRDA, evidenced by the enrichment of terms including signaling receptor activator activity (GO:0030546), signaling receptor regulator activity (GO:0030545), cytokine receptor binding (GO:0005126), small GTPase-mediated signal transduction (GO:0007264) and receptor ligand activity (GO:0048018). No single directional pattern dominated across all cell types, indicating that enrichment direction is context-dependent even when the same pathways persist.

**Figure 7:**
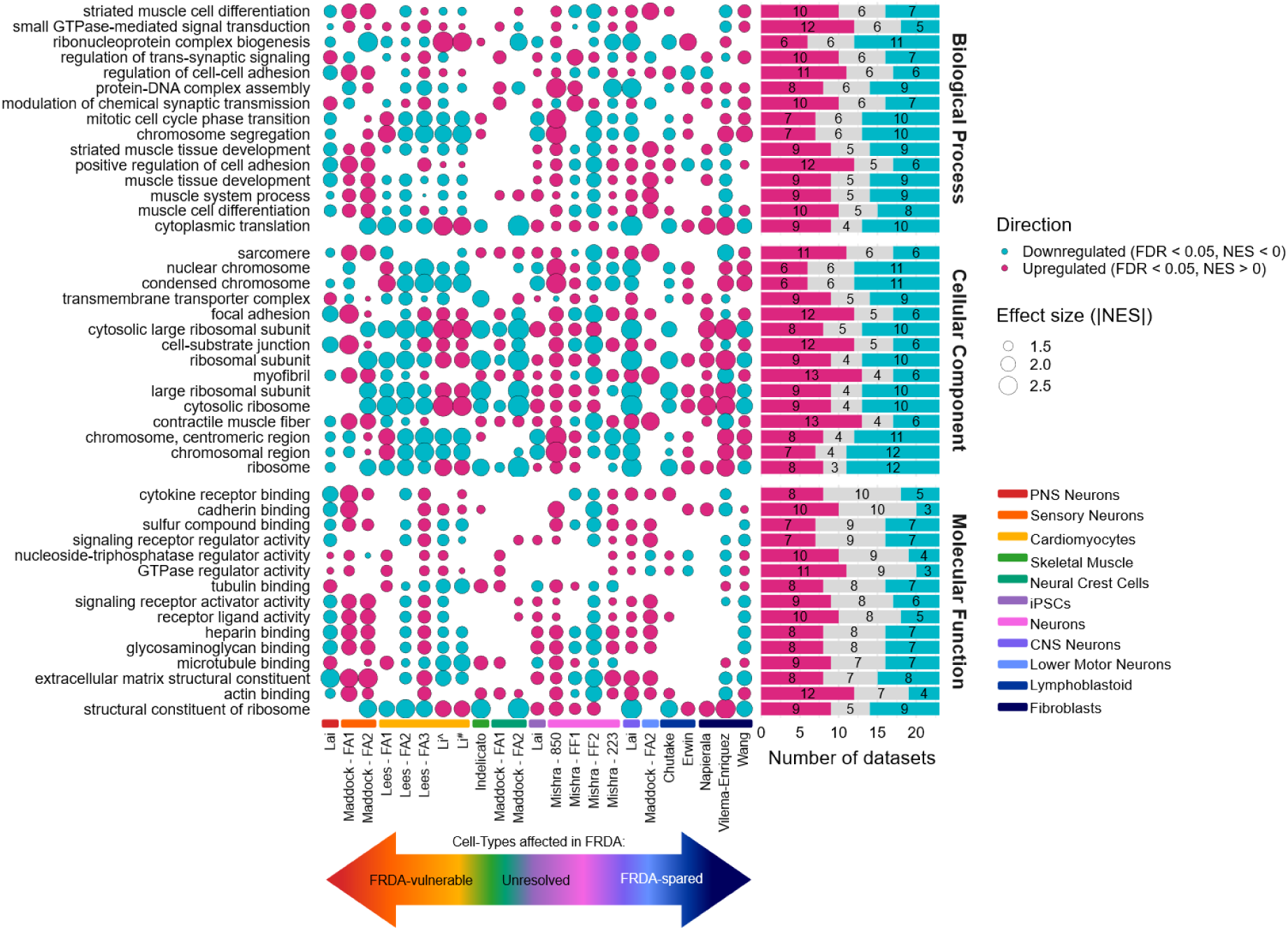
Recurrent translation, cytoskeletal and signaling process enrichment across FRDA datasets. Gene set enrichment analysis was performed independently for each FRDA versus control dataset using clusterProfiler across gene ontology biological process, cellular component and molecular function categories. Only gene sets containing 10 – 500 genes were included, and pathways with FDR < 0.05 were considered statistically significant. The top 15 terms for each gene ontology category are shown. The size of each point reflects absolute normalised enrichment score (|NES|) and colours indicate direction (pink: NES > 0, upregulated in FRDA; blue: NES < 0, downregulated in FRDA). Bar plots summarise the number of datasets enriched in each pathway.

### Stratified Analysis Reveals Distinct Cell-Type Dependent Dysregulation in FRDA

We next stratified the analyses by cell-type to examine top up and downregulated genes and pathways within each category, revealing distinct transcriptional patterns (Figure 8). The top 10 up and downregulated genes for each cell-type category are given in Figure 8A. The entire ranked gene lists for each cell type category are given in Supplementary File 2. Consistent with the global meta-analysis of the top genes in FRDA, stratification by cell type revealed similar genes dominated the top up and downregulated genes, for example *MEG3, MEG8, MEG9* and *MYH14* in sensory neurons. However overall, each cell type showed heterogeneous and context-specific gene changes, supporting the view that transcriptional dysregulation in FRDA is cell-type dependent.

**Figure 8:**
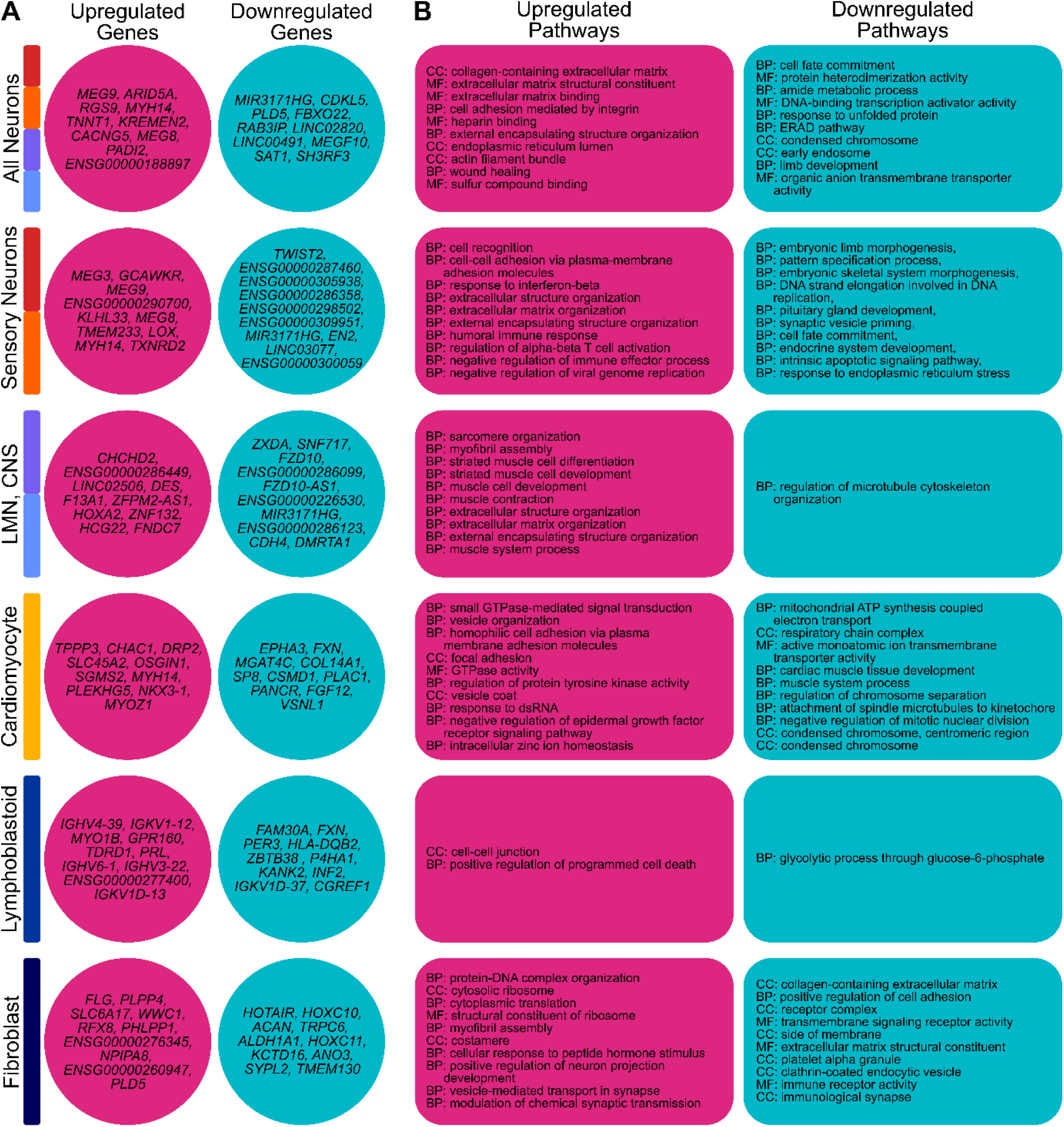
Cell-type specific transcriptional dysregulation in FRDA. A) Top 10 upregulated and downregulated genes identified within each cell-type category based on recurrence of differentially expression, directional consistency and median log_2_FC across FRDA versus control comparisons. B) Cell-type specific gene set enrichment analysis (GSEA) results, ranked by the number of datasets showing significant enrichment, directional consistency, and median normalised enrichment score. Redundant Gene Ontology (GO) terms were collapsed using semantic similarity-based reduction (rrvgo, similarity threshold = 0.7), with representative parent pathways shown for the top 10 positively and negatively enriched terms. Only pathways with a directions score > 0 are displayed. Complete ranked gene and pathway lists are provided in Supplementary Files 2 and 3.

To interpret how these recurrent gene level changes translate into broader biological processes, we next examined cell-type specific patterns of pathway enrichment within each category. All ranked pathways are given in Supplementary File 3. Sensory neurons were characterised by enrichment of developmental and patterning pathways (GO:0030326, GO:0007389, GO:0048704, GO:0021983, GO:0045165, GO:0035270), downregulation of synaptic vesicle dynamics (GO:0016082), apoptotic signaling (GO:0097193) and endoplasmic reticulum stress pathways (GO:0034976) in downregulated genes, suggesting dysregulation of developmental and stress-response processes. In CNS and LMNs, regulation of microtubule cytoskeleton organisation was enriched in downregulated genes (GO:0070507), pointing towards impaired cytoskeletal dynamics in these cells, rather than developmental processes seen in the sensory neurons. Downregulated pathways in cardiomyocytes were dominated by mitochondrial processes (GO:0042775, GO:0098803) including ATP synthesis, deficient cardiac muscle development (GO:0048738, GO:0003012) and chromosome function (GO:1905818, GO:0008608, GO:0045839, GO:0000779, GO:0000793). Lymphoblastoid cells exhibited suppression of glycolytic process through glucose-6-phosphate (GO:0061620) and fibroblasts displayed downregulation of pathways related to extracellular matrix organisation and cell-cell interactions (GO:0062023, GO:0045785, GO:0043235, GO:0004888, GO:0045334). Together, these findings suggest that transcriptional changes in non-neuronal cell types are characterised by reduced immune signaling, extracellular matrix dynamics, and cellular metabolism, consistent with a shared stress or secondary response rather than cell type–specific FRDA pathology.

Across cell types, upregulated pathways revealed both recurrent and context-specific transcriptional responses in FRDA. FRDA neuronal populations showed convergence on extracellular matrix organisation, cell adhesion and cytoskeletal remodelling, with all neurons enriched for integrin-mediated adhesion (GO:0033627), extracellular matrix components (GO:0062023, GO:0005201, GO:0050840), actin filament organisation (GO:0032432), and wound healing associated processes (GO:0042060), suggesting activation of structural and reparative programmes. Sensory neurons additionally exhibited upregulation of immune and interferon related pathways (GO:0035456, GO:0046634, GO:0002698, GO:0045071), indicating engagement of immune or stress responsive signaling alongside extracellular remodelling. In contrast, CNS and LMNs were dominated by upregulation of muscle and contractility related pathways (GO:0045214, GO:0030239, GO:0051146, GO:0055002, GO:0055001, GO:0006936, GO:0003012), consistent with altered cytoskeletal architecture rather than immune activation. Cardiomyocytes displayed a distinct profile characterised by increased small GTPase-mediated signaling and activity (GO:0007264, GO:0003924), vesicle organisation (GO:0016050), focal adhesion (GO:0005925), regulation of protein tyrosine kinase activity (GO:0061097) and zinc ion homeostasis (GO:0006882), reflecting altered intracellular signaling and trafficking. Non-neuronal cell types showed different dominant themes, with lymphoblastoid cells enriched for cell-cell junction (GO:0005911) and programmed cell death (GO:0043068), and fibroblasts marked by upregulated genes enriched in increased translational machinery (GO:0002181), myofibril anchoring (GO:0030239), and ribosome constituents (GO:0022626, GO:0003735). Together, these findings indicate that while structural and extracellular remodelling is a recurrent feature across FRDA cell types, the specific biological programmes engaged are strongly cell-type dependent.

### Treatment Responsiveness of Prioritised Transcriptomic Biomarkers

To evaluate whether consistently dysregulated genes in FRDA also respond to therapeutic intervention, we examined their behaviour in all available FRDA treatment datasets as a proof-of-concept assessment of their suitability as transcriptomic biomarkers. Across different treatment types, a subset of genes exhibited significant treatment-associated expression changes of the top 40 FRDA candidate biomarkers, with effects that either reversed or reinforced the FRDA versus control direction of differential expression (Figure 9). Directionally reversing responses were observed for multiple genes, including HDACi 109 and synTEF1 increasing *FXN* expression in iPSC-derived neurons and lymphoblastoid cells respectively. The antisense oligonucleotides S102 and S10L6 targeting *FXN*, and HDACi 109 also reversed the long intergenic non-coding RNA *LINC01515*, indicating that these disease-associated transcriptional signatures are pharmacologically modifiable. Other candidate biomarkers showed directionally concordant changes, suggesting reinforcement of FRDA-associated expression patterns, for example antisense oligonucleotides targeting *FXN* (S102, S10L6, S30) increasing *MEG9* expression in fibroblasts. Treatment responses varied across genes and experimental contexts, with effect sizes ranging from modest to pronounced, and no single intervention uniformly modulated all biomarkers, although the HDACi 109 affected the largest number of genes overall, consistent with its global epigenetic effects on transcription. Together, these results demonstrate that transcriptomic biomarkers are responsive in specific treatment contexts.

**Figure 9:**
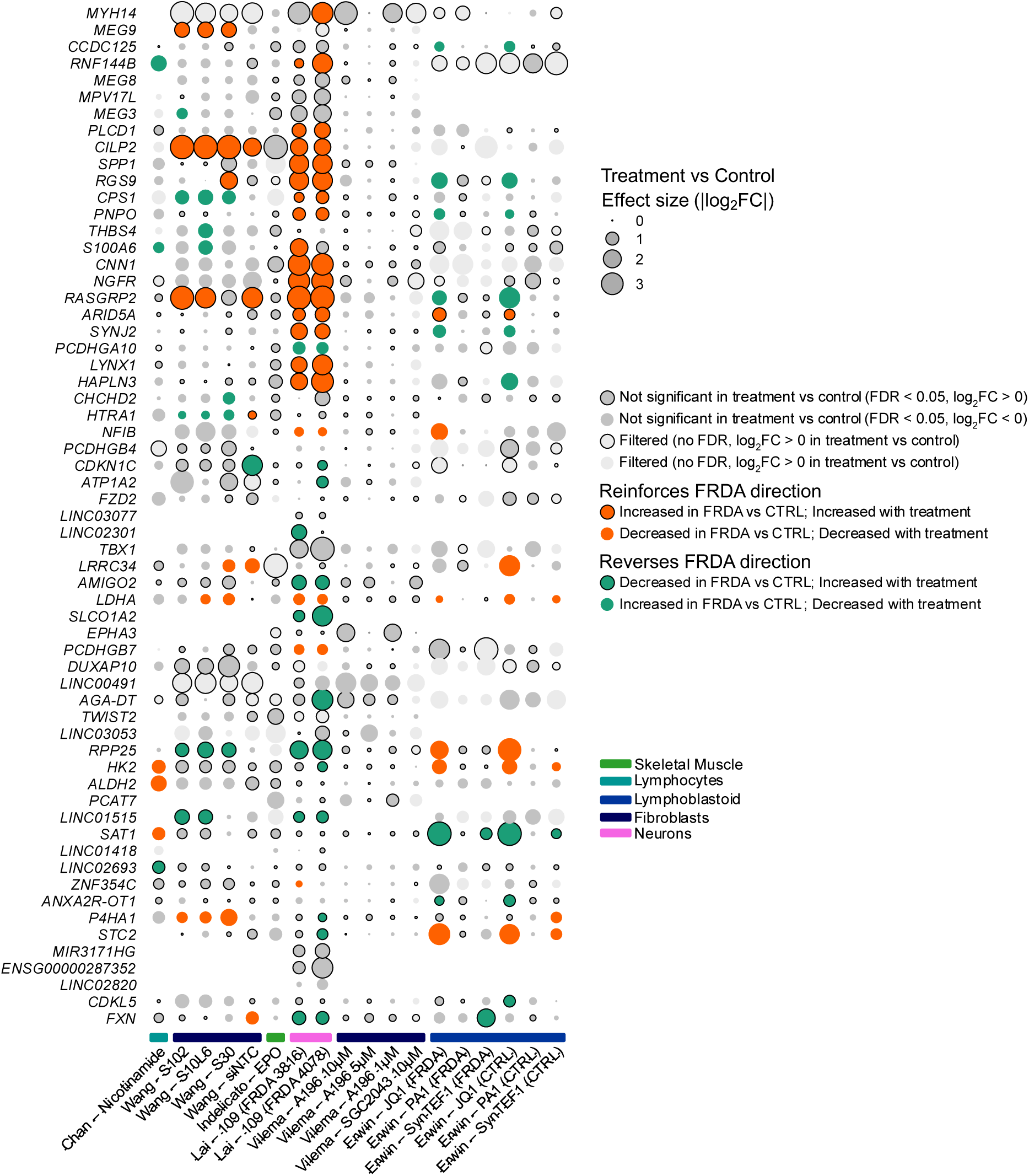
Therapeutic modulation of the 40 most consistently dysregulated FRDA-associated genes across independent FRDA treatment RNA-seq datasets. Publicly available FRDA treatment RNA-seq datasets spanning multiple cell types and therapeutic strategies were analysed, including nicotinamide treatment, antisense oligonucleotides (S102, S10L6, S30), erythropoietin (EPO), HDAC inhibitor (109), inhibitor of histone methyltransferase SUV20-H1 (A-196, 1 µM, 5 µM, 10 µM) and synthetic transcription elongation factors (syn-TEF1, JQ1, PA1). Treatment datasets were processed with the same pipelines used in the FRDA vs control datasets. Differential expression was computed for each treatment relative to a control within the same study (e.g., untreated control, vehicle control, siNTC). Point size reflects absolute log2 fold change (|log2FC|). Significant treatment-associated changes (FDR < 0.05) are coloured to indicate whether the treatment response reverses the original FRDA vs control direction (green; opposite to FRDA vs control) or reinforces it (orange; same direction as FRDA vs control). For non-significant genes (FDR ≥ 0.05) in treated vs control, points are coloured dark grey and have a black outline if the gene was expressed higher in treated compared to controls, or no outline if expressed lower in treated compared to controls.

### Limited Transcriptomic Dysregulation Among Canonical FRDA-Associated Markers

To determine whether previously reported FRDA-relevant molecular markers showed consistent gene-level changes, we examined a curated set of genes encoding proteins implicated in pathways disrupted in FRDA, including iron homeostasis, iron–sulfur cluster biogenesis, mitochondrial function and oxidative stress responses (54). Across the compiled datasets, most of these markers showed no significant difference in expression between FRDA and controls, including in highly affected cell types such as cardiomyocytes and sensory neurons (Figure 10). Aside from *FXN*, the only gene with a notable cross-dataset signal was *CAT* (catalase), which was significantly upregulated in FRDA in 8/23 and downregulated in 1/23 datasets. *CAT* expression appeared strongly donor-specific, with all FA1-derived cell types (iPSC-derived sensory neurons, neural crest cells, and cardiomyocytes) showing consistent upregulation. Overall, these findings indicate that many canonical FRDA-related markers may exhibit disease-associated changes primarily at the protein level rather than through alterations in steady-state mRNA abundance and are therefore unlikely to serve as robust transcript-level biomarkers.

**Figure 10.**
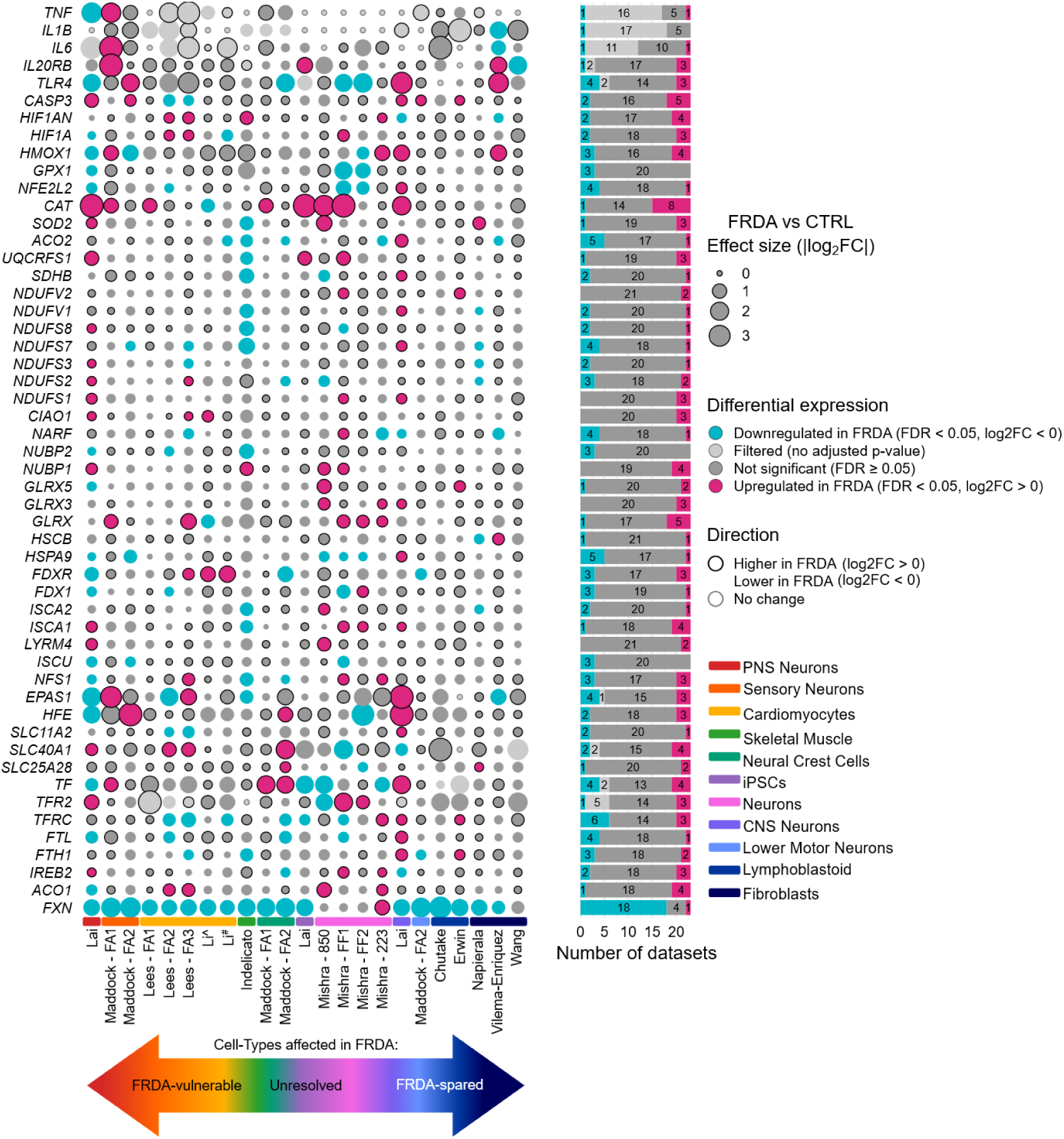
Limited gene-level changes among canonical FRDA-associated markers. Differential expression of a curated panel of FRDA-associated genes (iron-sulfur cluster biogenesis, mitochondrial function and oxidative stress responses) was assessed across all FRDA versus control RNA-seq datasets. Each point represents a dataset specific contrast for a given gene, with point size reflecting absolute log2 fold change (|log2FC|). Colour indicates differential expression status (pink: upregulated in FRDA, FDR < 0.05; blue: downregulated in FRDA, FDR < 0.05; grey: non-significant or filtered). Symbol outline denotes direction of change (higher in FRDA, lower in FRDA, or no change). Bar plots summarise the number of datasets showing upregulation, downregulation, or no significant change per gene.

## Discussion

Transcriptional profiling has been widely used to identify disease mechanisms of FRDA, and numerous RNA-seq studies have proposed candidate biomarkers beyond *FXN* based on their most pronounced DEGs. However, these candidates are rarely evaluated across independent FRDA datasets and diverse cell types, leaving their generalisability largely untested. Robust biomarkers, particularly those intended to determine therapeutic efficacy in clinical trials or *in vitro* drug screening, are unlikely to emerge from single RNA-seq studies alone. Moreover, the pronounced cell-type specific vulnerability of FRDA pathology, raises the possibility that transcriptional biomarkers identified in one cellular context may not translate to others. To address these limitations, we performed a systematic review and meta-analysis of human bulk RNA-sequencing datasets in FRDA, reprocessing all studies through a unified analysis pipeline to enable direct cross-study comparison. We analysed 23 independent datasets comprising 94 FRDA and 99 control samples across 10 distinct cell types, spanning both disease-relevant (cardiomyocytes, sensory neurons) and FRDA-spared cell types (fibroblasts, lymphoblastoid cells). This integrative approach enabled identification of transcriptional changes associated with FRDA across studies, while distinguishing those that are cell-type specific. By resolving both conserved and cell-type restricted transcriptional responses to frataxin deficiency, this framework highlights candidate genes and pathways that may help explain how ubiquitous frataxin loss gives rise to selective tissue vulnerability. Given the limited availability of robust molecular biomarkers in FRDA, we identified a panel of candidate transcriptional biomarkers with potential utility in both *in vitro* drug screening and clinical trial settings. All candidate biomarkers, datasets and results are accessible through the open-access FRDA Transcriptomic Atlas (37).

### Beyond Frataxin: Recurrent Transcriptomic Dysregulation in FRDA

Based on the most consistently upregulated and downregulated genes across datasets, we defined a panel of candidate transcriptomic biomarkers with potential utility for therapeutic screening *in vitro* and as efficacy readouts in clinical trials (Table 4). Several top-ranked genes including *MYH14*, *MEG9* and *CDKL5* were not highlighted as key findings in the original individual RNA-seq studies yet emerged as consistently dysregulated when assessed across multiple independent datasets, highlighting the value of cross-study recurrence analyses. These candidate biomarkers have the potential to bridge *in vitro* and clinical readouts and to expand the biomarker “toolbox” beyond frataxin and mitochondrial function.

*FXN* remained the most consistently downregulated gene across datasets, reinforcing its position as the primary transcriptomic biomarker for FRDA. Notably, a few datasets did not show significant *FXN* reduction despite clear disease phenotypes. These discrepancies likely reflect a combination of limited statistical power and model-specific effects including differences in genetic backgrounds and isogenic control systems. Importantly, these datasets showed significant frataxin reduction at the protein level, highlighting that *FXN* mRNA abundance does not necessarily predict protein levels. Additionally, pathogenic effects of the GAA expansion independent of *FXN* depletion may contribute to disease-associated transcriptional changes (55,56), supporting inclusion of these datasets within the meta-analysis. Other biological factors may also contribute to a lack of *FXN* reduction in some datasets. For example, interruptions within the expanded GAA tract (i.e., non-pure GAA repeats) have been reported to partially restore *FXN* transcription and delay disease onset (57–59). However, detailed characterisation of GAA repeat structure, including repeat interruptions, was not routinely performed when many of these RNA-seq datasets were generated, and therefore this information is unavailable for most datasets included in this meta-analysis.

Apart from *FXN*, the most recurrent significantly downregulated gene across FRDA was *CDKL5*. *CDKL5* downregulation was not consistent across cell types or donor backgrounds, suggesting potential donor-specific effects. *CDKL5* encodes a serine/threonine protein kinase involved in microtubule and actin dynamics, contributing to axon growth in neurons, synaptogenesis, dendritic arborization, neuronal development and cell proliferation (60–67). Loss of CDKL5 causes CDKL5 deficiency disorder, a severe neurodevelopmental condition characterised by childhood-onset epilepsy, hypotonia, motor and cognitive disabilities (68–70). Beyond its role in neuronal function, CDKL5 regulates mitochondrial function. CDKL5 deficiency is associated with reduced oxidative phosphorylation complex subunits, mitochondrial trafficking and impaired complex I and IV activity in iPSC-derived neurons (71). Given that mitochondrial dysfunction is central to FRDA pathogenesis, reduced *CDKL5* expression may further exacerbate mitochondrial stress by compromising oxidative phosphorylation or intracellular transport. CDKL5 deficiency has also been associated with reduced NRF2 levels and impaired NRF2-mediated oxidative stress responses in fibroblasts (72), mirroring defects observed in FRDA. Suppressed NRF2 signaling is a recognised component of FRDA pathology and underlies the mechanism of action of Omaveloxolone, the only approved disease-modifying treatment for individuals with FRDA ≥ 16 years. Whether CDKL5 directly modulates NRF2 activity in FRDA remains to be determined. Notably, *CDKL5* expression did not respond to any therapeutic interventions analysed in this study and showed variable downregulation across cell types and donor backgrounds, suggesting donor-specific regulation that may influence therapeutic responsiveness. As CDKL5 restoration improves phenotypes in experimental models (71), determining whether its downregulation is reversible in FRDA phenotypic subgroups may have therapeutic relevance.

The most consistently upregulated gene across FRDA datasets was *MYH14*, encoding the non-muscle myosin II-C (NMIIC) motor protein. NMIIC belongs to the non-muscle myosin II family (NMII) alongside *MYH9* (NMIIA) and *MYH10* (NMIIB), which share roles in contractility and regulating cell morphology through the cytoskeletal network. In contrast to the broadly expressed NMIIA and NMIIB isoforms, NMIIC shows more restricted expression and remains comparatively understudied (73). Notably, NMIIC is enriched in the cerebellum, skeletal muscle and heart (74), tissues prominently affected in FRDA, suggesting that *MYH14* upregulation in FRDA may be particularly relevant within disease-vulnerable lineages. Further, NMIIC overexpression promotes mitochondrial fission and fragmentation (75). Consistent with this, increased mitochondrial fission has been reported across multiple FRDA models, including iPSC-derived cardiomyocytes (49), *FXN* knockdown fibroblasts, patient fibroblasts (19) and *FXN* knockout mice exhibiting elevated Drp1 expression *in vivo* (76). These observations raise the possibility that *MYH14* upregulation contributes to altered mitochondrial dynamics in FRDA. Importantly, NMIIC motor activity is ATP-dependent, and ATP depletion is reported in multiple FRDA models (17–21,77–80), indicating that increased *MYH14* expression may not translate into increased NMIIC activity under energetic stress. Accordingly, gene expression alone is insufficient to infer functional consequences, and direct experimental interrogation of NMIIC activity will be essential to determine whether *MYH14* upregulation is adaptive, maladaptive, or epiphenomenal. Interestingly, the HDACi 109 further increased *MYH14* expression in treated iPSC-derived neurons compared to unaffected controls, suggesting epigenetic responsiveness. Establishing the functional significance of *MYH14* upregulation will be important for evaluating its potential utility as a FRDA biomarker in therapeutic screening contexts.

The next most consistently upregulated gene in FRDA was the long non-coding RNA *MEG9*. *MEG9* resides in the imprinted *DLK1-DIO3* locus, a genomic region that also includes the lncRNAs *MEG8* and *MEG3*. Both *MEG8* and *MEG3* were also frequently significantly upregulated, suggesting locus-level dysregulation rather than isolated gene-specific effects. lncRNAs within the *DLK1-DIO3* locus are implicated in epigenetic regulation (81,82), cellular stress responses (82,83) and vascular and neuronal homeostasis (83–85), although individual lncRNAs at this locus exert divergent and sometimes opposing functions. For example, *MEG3* promotes apoptosis (49,81,86,87), whereas *MEG9* overexpression promotes survival under stress in endothelial cells (83). Fralie-Bethencourt et al., 2022 also observed that *MEG9* and *MEG3* respond in opposite directions to DNA damage(83). Particularly, Lees et al., 2025, were the first to speculate that *MEG3* upregulation has a pro-apoptotic role in FRDA iPSC-derived cardiomyocytes (49). Together, this indicates that upregulated *MEG3* may reflect a maladaptive stress response, whereas increased *MEG9* may be a compensatory or protective response. However, *MEG9* is comparatively understudied relative to *MEG3* and *MEG8*, and its role in FRDA requires clarification. Notably, treatment of FRDA fibroblasts with the *FXN*-targeting antisense oligonucleotides (S10, S10_L6) significantly upregulated *MEG9*, indicating responsiveness to *FXN*-restoring therapy (40), whereas *MEG8* showed no modulation across treatment datasets. This differential responsiveness supports non-uniform regulatory control and functional divergence within the *DLK1-DIO3* locus.

*MEG9*, *MEG8*, and *MEG3* exhibited cell-type specific upregulation, with increased expression in FRDA-relevant cell types including PNS sensory neurons and cardiomyocytes, but not in skeletal muscle, CNS neurons, lymphoblastoid cells and fibroblasts, supporting a role for these lncRNAs in FRDA cell-type vulnerability. Upregulation of *MEG9*, *MEG8*, and *MEG3* in iPSCs and a neural crest cell dataset suggests that dysregulation of the *DLK1-DIO3* locus may arise early during differentiation and predispose specific lineages to disease-associated vulnerability. However, the absence of consistent dysregulation in several iPSC-derived lineages, including CNS neurons and lower motor neurons, indicates that this perturbation is not universally maintained, but instead becomes functionally relevant in selectively vulnerable FRDA cell types.

### Considerations for Biomarker Use in FRDA: Cell-Type Relevance

Many of the strongest candidate biomarkers identified were consistently dysregulated only in disease-relevant cell types. This has important implications for clinical trial design where therapeutic efficacy is frequently assessed using accessible tissues such as blood or fibroblasts that may not reflect core disease biology. Our findings emphasise the need to align biomarker selection with tissue relevance and caution against overinterpreting transcriptional changes measured in cell types that are not directly affected in FRDA. *FXN* was the only gene consistently differentially expressed across both affected and unaffected cell types, supporting its utility as a reliable transcript-level biomarker in peripheral samples. *RNF144B* also emerged as a promising exception, displaying significant upregulation in peripheral blood in 10/23 of the datasets analysed in this study, suggesting that certain biomarkers may link peripheral readouts with disease-relevant molecular pathology and thus have greater translational potential. This distinction is critical for clinical feasibility, where access to disease-relevant tissues is often limited. Given their relevance to FRDA pathology, skeletal muscle biopsies may capture treatment-associated transcriptional alterations that are not readily detected in whole blood or fibroblasts. For example, *CHODL* and *S100A6* were upregulated in both skeletal muscle and sensory neurons, suggesting muscle expression may serve as an indirect readout of sensory neuron-related molecular changes. Among candidate biomarkers, *MEG9* showed promising but heterogeneous dysregulation in skeletal muscle, with a subset of apparent FRDA non-responders. In the control group *MEG9* was largely undetectable (0 – 0.48 TPM) but highly expressed in 4/7 FRDA samples (0 – 5.68 TPM). This apparent “all-or-nothing” expression pattern may make *MEG9* a particularly sensitive biomarker for use. Transcriptomic profiling may therefore complement longitudinal clinical outcomes, such as clinical rating scales, by capturing a real-time molecular snapshot, capable of detecting early therapeutic responses that precede observable clinical improvement. Although the clinical implications of *MEG9* dysregulation is yet to be established. RNA-seq also enables unbiased, transcriptome-wide assessment of cellular state, capturing molecular dysfunction across a range of pathways that may be missed by targeted assays or readouts.

While this meta-analysis prioritises gene biomarkers dysregulated across independent FRDA datasets, gene level association alone does not establish functional relevance. Future studies must determine whether these expression changes are compensatory responses to frataxin deficiency or contributors to FRDA pathology to be useful biomarkers. Although functional involvement is not required for a biomarker to be clinically useful, such studies will help distinguish markers that track disease state from those that may represent mechanistic drivers or actionable therapeutic targets.

### Considerations for Using Classical FRDA-related Markers at the Transcript Level

Across all cell-types, canonical ‘FRDA marker’ genes showed little significant differential expression between individuals with FRDA and controls. These genes encode proteins involved in iron homeostasis, Fe-S cluster biogenesis, oxidative phosphorylation, mitochondrial function and oxidative stress resolution; processes that are well established to be disrupted in FRDA. The absence of transcriptional changes suggests that these defects arise primarily through post-transcriptional mechanisms and altered protein function downstream of frataxin deficiency, rather than coordinated shifts in steady-state mRNA abundance. This finding indicates that traditional FRDA marker genes are poorly suited as RNA-level biomarkers. Instead, transcript classes consistently dysregulated at the RNA level, particularly long non-coding RNAs identified in the biomarker panel (Table 4), may provide informative readouts of disease-associated transcriptional reprogramming. A complementary meta-analysis of available FRDA proteomic datasets will therefore be necessary for identifying protein-level biomarkers that better capture disease-associated dysfunction not evident at the transcript level.

### Cell-Type Resolved Pathway Dysregulation in FRDA

Gene set enrichment analysis further supported the absence of typical FRDA-associated pathways changes such as iron and mitochondrial dysfunction, at the transcript level. Instead, enrichment was dominated by ribosomal/translation, cytoskeletal, and signaling pathways, suggesting that frataxin deficiency propagates downstream effects into broader cellular homeostasis pathways detectable transcriptionally. Mitochondrial dysfunction and oxidative stress are well known to suppress global translation through stress-response signaling (88), providing a plausible mechanistic link between frataxin loss and the observed translation-related signatures. Consistent with this, altered expression of transcriptional and translational machinery has been reported across multiple FRDA models, including reduced translation pathways in FRDA peripheral blood mononuclear cells, which correlated with disease severity (44), and decreased protein synthesis factors and ribosomal proteins in FRDA fibroblasts (50), B-lymphocytes (89) and skeletal muscle (90).

Cytoskeletal pathways were also recurrently enriched across datasets. Cytoskeletal dysfunction has been proposed as an underappreciated disease mechanism in FRDA (91). In FRDA fibroblasts, actin underwent glutathionylation, a redox-sensitive post-translational modification that impaired actin polymerisation and hence microfilament organization and cytoskeletal dynamics (92). Frataxin silenced motor neurons also showed an increase in glutathione bound to the α-tubulin component of the cytoskeleton, impacting microtubule organisation, microtubule dynamics and neurite extension (93). Thereby, cytoskeletal abnormalities may extend to the cell types measured in this study. This is further supported at the gene level by dysregulation of *MYH14*, responsible for interacting with the cytoskeleton for contraction, cytokinesis and initiating cell adhesion to the extracellular matrix (73). Collectively, these observations motivate targeted functional validation studies to determine whether translational and cytoskeletal dysregulation represent robust, quantifiable biomarker axes in FRDA and potential targets for therapeutic development.

Although the global meta-analysis did not reveal uniformly directionally consistent pathway changes across all datasets, stratification by cell type resolved distinct and biologically coherent transcriptional pathways. This finding underscores the importance of cellular context in interpreting FRDA transcriptomic data. Separation into neuronal, cardiac, and non-neuronal categories revealed convergent yet cell-type specific pathway dysregulation aligned with physiological vulnerabilities of each cell type. For example, mitochondrial and contractile deficits were predominant in cardiomyocytes, alterations in synaptic dynamics in sensory neurons and upregulated ribosome and translational components in fibroblasts. These findings indicate that the absence of universal directional enrichment across all FRDA datasets does not reflect a lack of biological signal, but rather the presence of highly cell-type restricted transcriptional responses, potentially representing distinct pathophysiological or adaptive processes in response to frataxin deficiency.

Sensory neuron and cardiomyocyte categories showed enrichment patterns consistent with disruption of developmental and morphogenetic programmes, including downregulated pattern specification, cell fate commitment and morphogenesis in sensory neurons, and reduced cardiac muscle development and muscle system processes in cardiomyocytes. Such pathways were not prominent in fibroblast, lymphoblastoid, or relatively spared neuronal cells such as lower motor neurons and CNS neurons. These observations support an emerging hypothesis of FRDA as both a developmental and neurodegenerative disorder, in which early lineage perturbations may predispose vulnerable cell types to impaired maturation, contributing to later functional decline under lifelong frataxin deficiency. This finding aligns with further transcriptomic evidence showing that developing iPSC-derived sensory neurons have more DEGs than their more mature counterparts, suggesting heightened vulnerability during development of sensory neurons (54). Pathological disruption at the dorsal root entry zone, where astroglial tissue breaches the CNS-PNS boundary in FRDA, further supports a developmental component to FRDA pathology (94). As this boundary is established by neural crest cell-derived boundary cap cells during development, its alteration likely arises embryonically rather than through degeneration alone. Together, this raises the possibility that frataxin restoration in adulthood may be insufficient to address earlier developmental defects.

More broadly, the cell-type specificity of dysfunctional pathways in FRDA has important implications for biomarker interpretation, therapeutic targeting, and study design. Biomarkers identified in one cellular context may not generalise across tissues, and conclusions drawn from bulk or mixed cell datasets risk misrepresenting disease biology. These findings suggest FRDA represents a spectrum of context-dependent molecular adaptations rather than a single transcriptomic state that may underpin the selective vulnerability of certain cell-types in FRDA.

### Considerations for Reproducibility and Study Design in future FRDA RNA-seq studies

As part of this systematic review, we developed a quality assessment framework to evaluate experimental design and reporting across FRDA RNA-sequencing studies and areas for improvement. This framework comprised 21 categories, scored on a three-point scale (low, medium, high), and was intended both to benchmark existing datasets and to provide a practical checklist for future FRDA transcriptomic studies. One of the low-scoring categories concerned the clear distinction between biological and technical replication, particularly within iPSC-derived cohorts. Owing to the cost and logistical complexity of iPSC generation, many studies were constrained to using cells with limited genetic diversity and defined biological replicates as independent differentiations rather than genetically distinct donors. When multiple FRDA cell lines were included, reporting was often insufficient to determine whether differentiations were analysed per donor or pooled across donors. In several cases, repeated differentiations from a single donor appeared to be combined with differentiations from other donors and analysed as a single FRDA group, effectively inflating replicate numbers instead of representing true biological independence. Such practices inflate effect sizes as the variability between differentiations is typically lower than between different genotypes unless explicitly modelled. Differentiation replicates should therefore assess technical variability, whereas biological replication requires multiple genetically distinct donors analysed independently to identify shared molecular signatures across genotypes. Clear reporting of replicate definitions and analytical strategy is essential to avoid pseudo-replication and enable accurate interpretation and cross-study comparability.

Beyond replication, sample size was another low-scoring category. Five studies included only a single donor with FRDA, and only two studies, skeletal muscle and fibroblast cohorts incorporated more than three independent donors with FRDA. Although this meta-analysis included 94 FRDA samples, these represented just 35 unique donor backgrounds, with 18 originating from a single fibroblast dataset (Napierala), reflecting extensive reuse of a small number of well-characterised cell lines. For example, Coriell Institute cell line GM03816 was used across five datasets and differentiated into several cellular contexts (fibroblasts, iPSCs, iPSC-derived neurons, iPSC-derived CNS neurons, iPSC-derived PNS neurons). While these lines have been invaluable for dissecting cell-type specific effects within a controlled genetic background, their repeated use limits assessment of whether identified transcriptional biomarkers generalise across patients or reflect donor-specific effects. This limitation is particularly relevant in FRDA where GAA repeat length correlates with disease severity and age of onset (95). Larger and more genetically diverse cohorts will be required to assess whether candidate biomarkers identified here correlate with GAA expansion size.

This meta-analysis highlights a major limitation of the current FRDA transcriptomic landscape – the absence of postmortem RNA-seq data and a strong bias toward iPSC-derived models, which comprised most included datasets. While iPSC models enable study of otherwise inaccessible human cell types, they typically represent developmentally immature states that lack the lifelong physiological stress and disease progression experienced *in vivo*. As a result, it remains unclear whether the candidate biomarkers identified here persist in late-stage disease or instead reflect early molecular responses to frataxin deficiency that drive disease pathology. Beyond these tissues, several clinically relevant cell populations remain unexplored, including pancreatic β cells, given the higher prevalence of diabetes in FRDA, retinal cell types underlying visual dysfunction, and glial populations of the central and peripheral nervous systems. While differentiation protocols for these cell types exist, comprehensive transcriptomic characterisation and direct comparison with controls are lacking, and 3D model systems such as cerebellar organoids remain underutilised. Expanding RNA-seq profiling to these disease relevant tissues and models will be critical for defining stage and tissue specific molecular signatures and determining whether candidate biomarkers identified in iPSC-based systems generalise to clinically relevant contexts.

### Limitations of Transcriptomic Meta-Analyses: Considerations for Future Studies

Other limitations of this study primarily relate to scope and data availability. Datasets lacking peer-reviewed publication or GEO deposition may have been missed, potentially biasing inclusion toward more recent or better-resourced studies. This review also focused exclusively on bulk RNA-seq to enable standardised re-analysis across datasets, thereby excluding single-cell and spatial transcriptomic studies increasingly used to resolve cell-type specific pathology. A substantial body of FRDA transcriptomic profiling based on microarray (16,44,96–100) and miRNA (101,102) profiling was also excluded to preserve analytical consistency, which reduced the total evidence base. Likewise, other model systems, including yeast and *Drosophila* models, *FXN* knockdown or overexpression systems, and FRDA mouse models (e.g. KIKO and YG8R), were excluded, limiting assessment of whether the identified transcriptomic signatures are conserved across species or are specific to human disease. Together, these constraints restrict the breadth and generalisability of the evidence base and highlight the need for integrative comparisons across sequencing types and non-human models to identify recurrent dysregulated genes in FRDA.

No single gene was consistently dysregulated across all datasets, likely reflecting both biological and technical variability. Key technical contributors include sequencing depth and library preparation methods. Studies sequenced at low depth may have incomplete transcriptome coverage, causing low-abundance genes to appear absent or non-significant, not because they are unregulated, but because they were insufficiently sampled. This limitation is particularly relevant for the Indelicato and Erwin cohorts, sequenced at ∼ 4-5 million reads per sample, which is below the recommended 30 million reads required to provide coverage of most genes (103). At this depth, transcriptome coverage is restricted to the most highly expressed genes, increasing the likelihood that low-abundance transcripts are not detected or only stochastically observed. Library preparation methods further contribute to inter-study heterogeneity. Poly(A) enrichment selectively captures mRNA with poly(A) tails and under-represents non-polyadenylated transcripts including many long non-coding RNAs, miRNAs and rRNAs. Consequently, studies using poly(A) enrichment (Chutake (LCL), Erwin (LCL), Indelicato (skeletal muscle), Li (cardiomyocytes), Mishra (neurons), Vilema-Enriquez (fibroblasts), Wang (fibroblasts)) may fail to detect dysregulation simply because such transcripts were not ascertained. Given the prominent dysregulation of long non-coding RNAs identified here (e.g. *MEG9*, *MEG8*, *MEG3*), future studies would benefit from prioritising rRNA depletion preparations over poly(A) enrichment to achieve more transcriptome coverage. In addition, variability in sequencing depth and library preparation constrained isoform-level analysis at the meta-analytic level, as several cohorts lacked sufficient read depth and splice junction coverage for reliable transcript resolution.

### The FRDA Transcriptomic Atlas: A One-Stop Platform for FRDA Transcriptomic Exploration

The FRDA Transcriptomic Atlas was developed to address a major limitation in individual RNA-seq studies: although many transcriptomic studies exist, their results are difficult to compare directly across models and laboratories. The FRDA Transcriptomic Atlas allows researchers and clinicians to explore the large amount of FRDA sequencing data that exists, enabling them to not only visualise the results of this meta-analysis, but also to find their favourite genes and pathways and create their own hypotheses based on the available data. Instructions on how to use the FRDA Transcriptomic Atlas are here: (37). We aim to maintain the Atlas as an up-to-date resource by incorporating newly available bulk RNA-sequencing datasets, ensuring its long-term relevance and refinement of the FRDA transcriptomic landscape to support biomarker discovery, therapeutic target identification, and cell-type specific treatment strategies.

## Conclusion

In this study we processed all available human bulk RNA-seq datasets in FRDA to generate a harmonised cross-study transcriptomic analysis and atlas. We identified a panel of candidate transcriptional biomarkers in FRDA, including multiple genes that exhibited dysregulation only in FRDA affected cell types. This candidate biomarker panel updates previously adopted FRDA gene sets in terms of prevalence of differentially expressed genes across studies. While *FXN* remains the most consistently dysregulated gene and the central hallmark of FRDA, our meta-analysis reveals additional convergent transcriptional signatures, including dysregulation of the *DLK1-DIO3* locus (*MEG9*, *MEG8*, *MEG3*), consistent downregulation of *CDKL5* and upregulation of *MYH14*, highlighting novel molecular pathways implicated in FRDA. The pronounced cell-type specificity of these signals highlights the importance of context-aware biomarkers for accurately capturing FRDA pathophysiology and for informing therapeutic development. To support this, we created the FRDA Transcriptomic Atlas, an open resource that enables examination of gene and pathway level dysregulation across studies and cell types in FRDA, facilitating hypothesis generation, biomarker prioritisation, and translational target selection. Collectively, this work refines the molecular landscape of FRDA and provides a scalable platform to accelerate biomarker-driven research and therapeutic strategies aimed at improving outcomes for individuals with FRDA.

## Abbreviations

ATP: Adenosine triphosphate
CDD: CDKL5 deficiency disorder
CNS: Central nervous system neuron
DEGs: Differentially expressed genes
DOI: Digital Object Identifier
FC: Fold-change
FDR: False discovery rate
FRDA: Friedreich ataxia
GAA: Guanine-adenine-adenine
GEO: Gene expression omnibus
GO: Gene ontology
GSEA: Gene set enrichment analysis
HDACi: Histone deacetylase inhibitor
iPSC: Induced pluripotent stem cells
lincRNA: Long intergenic non-coding ribonucleic acid
LMN: Lower motor neuron
lncRNA: Long non-coding ribonucleic acid
NES: Normalised Enrichment Score
PCA: Principal component analysis
PNS: Peripheral nervous system
PRISMA: Preferred Reporting Items for Systematic Reviews and Meta-Analyses
rhuEPO: recombinant human Erythropoietin
RIN: RNA Integrity Number
RNA: Ribonucleic acid
RNA-seq: RNA sequencing
rRNA: Ribosomal ribonucleic acid
RT-qPCR: Reverse transcriptase quantitative polymerase chain reaction
SD: Standard deviation
Syn-TEF1: Synthetic transcription elongation factor 1
TPM: Transcripts per million

## Declarations

### Ethics Approval

New datasets generated in this study were covered by Human Ethics approved by the University of Wollongong Human Research Ethics Committee (HREC 202/451) and Biosafety Approval (GT19/08).

### Consent for publication

Not applicable.

### Data Availability

New datasets generated in this study (FRDA and isogenic corrected iPSC-derived sensory neurons, neural crest cells and lower motor neurons) are available on the gene expression omnibus (GEO) with accession ID: GSE319887 (https://www.ncbi.nlm.nih.gov/geo/query/acc.cgi?acc=GSE319887). Other publicly available datasets used in this study with GEO IDs are detailed in Table 2.

### Authors’ contributions

**Conceptualization:** MLM, Mdott; **Investigation**: MLM, SM, JGL, CD, AD, AH; **Methodology**: MLM, SM, JGL; **Formal Analysis**: MLM; **Software and FRDA Transcriptomic Atlas development:** MLM; **Data Curation**: MLM; **Validation:** MLM, SM, AD, CK, AH, LAC, MDel, RKF-U, ANA, CD, JGL, JSN, MN, Mdott; **Resources**: MDott, SIB, JMG, MP, JGL, SYL, ES, JSN, MN; **Visualisation**: MLM; **Supervision:** Mdott, AH, RKF-U, ANA; **Project Administration**: Mdott; **Funding Acquisition**: Mdott, SYL, SIB, JMG, JGL, MP, JSN, MN; **Writing – original draft:** MLM; **Writing – review and editing:** All authors. All authors read and approved the final manuscript.

## Acknowledgements

This research was undertaken with the assistance of resources and services from the National Computational Infrastructure (NCI Australia), including the Gadi high-performance computing system, an NCRIS-enabled capability supported by the Australian Government. We would like to thank all FRDA researchers who generated and made their RNA-seq datasets publicly available. We also thank the groups who generously shared unpublished or non-public datasets, which substantially strengthened the breadth of this meta-analysis. This work was made possible by the commitment of the FRDA research community to data sharing. Marnie L. Maddock and Chloe K. Kennedy were supported by an Australian Government Research Training Program Scholarship. Amy J Hulme was supported by a FARA postdoctoral fellowship. Jarmon G. Lees was supported by a Future Leader Fellowship (108435-2024_FLF) from the National Heart Foundation of Australia.

