## Supplementary File 1 for "Friedreich ataxia transcriptomic dysregulation and identification of cell type-specific biomarkers: A systematic review and meta-analysis"

### Friedreich's ataxia transcriptomic dysregulation and identification of cell type-specific biomarkers: A systematic review and meta-analysis

Marnie L. Maddock[1]†, Sara Miellet[1], Anjila Dongol[1], Amy J. Hulme[1], Chloe K. Kennedy[1], Louise A. Corben[2,3,4], Rocio K. Finol-Urdaneta[1], Alberto Nettel-Aguirre[5,6], Chiara Dionisi[7], Martin B. Delatycki[8], Joel M. Gottesfeld[9], Massimo Pandolfo[10], Elisabetta Soragni[11], Sanjay I. Bidichandani[12], Jarmon G. Lees[13,14,15], Shiang Y. Lim[13,14,15,16], Jill S. Napierala[17], Marek Napierala[17], Mirella Dottori[1]†

† Co-corresponding authors:

Marnie L. Maddock:

Mirella Dottori:

1. Molecular Horizons, School of Medical and Indigenous Health Sciences, Faculty of Science, Medicine and Health, University of Wollongong, Wollongong, New South Wales, Australia
2. Murdoch Children's Research Institute, Parkville, Victoria, Australia
3. Department of Paediatrics, University of Melbourne, Parkville, Victoria, Australia
4. Turner Institute of Brain and Mental Health, Monash University, Victoria, Australia
5. School of Social Sciences, University of Wollongong, Wollongong, New South Wales, Australia
6. Department of Pediatrics, University of Calgary, Alberta, Canada
7. School of Immunology and Microbial Sciences, King's College London, London, UK
8. Bruce Lefroy Centre, Murdoch Children's Research Institute, Victoria, Australia
9. Scripps Research Institute, La Jolla, California, USA
10. Department of Neurology and Neurosurgery, McGill University, Montreal, Quebec, Canada
11. Friedreich's Ataxia Research Alliance, USA
12. Department of Pediatrics, University of Oklahoma College of Medicine, Oklahoma City, Oklahoma, USA
13. St Vincent's Institute of Medical Research, Victoria, Australia
14. Department of Medicine and Surgery, University of Melbourne, Victoria, Australia
15. Drug Discovery Biology, Monash Institute of Pharmaceutical Sciences, Monash University Parkville, Victoria, Australia
16. National Heart Research Institute Singapore, National Heart Centre, Singapore
17. Department of Neurology, Peter O'Donnell Jr. Brain Institute, University of Texas Southwestern Medical Center, Dallas, Texas, USA

Supplementary Table 1: PRISMA Checklist

| Section and Topic | Item # | Checklist item | Location where item is reported |
|---|---|---|---|
| **TITLE** | | | |
| Title | 1 | Identify the report as a systematic review. | Title. Methods: Systematic Review |
| **ABSTRACT** | | | |
| Abstract | 2 | See the PRISMA 2020 for Abstracts checklist. | Abstract |
| **INTRODUCTION** | | | |
| Rationale | 3 | Describe the rationale for the review in the context of existing knowledge. | Introduction |
| Objectives | 4 | Provide an explicit statement of the objective(s) or question(s) the review addresses. | Introduction p7 |
| **METHODS** | | | |
| Eligibility criteria | 5 | Specify the inclusion and exclusion criteria for the review and how studies were grouped for the syntheses. | Methods: Study Selection |
| Information sources | 6 | Specify all databases, registers, websites, organisations, reference lists and other sources searched or consulted to identify studies. Specify the date when each source was last searched or consulted. | Methods: Search Strategy |
| Search strategy | 7 | Present the full search strategies for all databases, registers and websites, including any filters and limits used. | Supplementary Table 2. |
| Selection process | 8 | Specify the methods used to decide whether a study met the inclusion criteria of the review, including how many reviewers screened each record and each report retrieved, whether they worked independently, and if applicable, details of automation tools used in the process. | Methods: Study Selection, Search Strategy |
| Data collection process | 9 | Specify the methods used to collect data from reports, including how many reviewers collected data from each report, whether they worked independently, any processes for obtaining or confirming data from study investigators, and if applicable, details of automation tools used in the process. | Methods: Data Extraction |
| Data items | 10a | List and define all outcomes for which data were sought. Specify whether all results that were compatible with each outcome domain in each study were sought (e.g. for all measures, time points, analyses), and if not, the methods used to decide which results to collect. | Methods: Data Extraction |
| | 10b | List and define all other variables for which data were sought (e.g. participant and intervention characteristics, funding sources). Describe any assumptions made about any missing or unclear information. | Methods: Data Extraction |
| Study risk of bias assessment | 11 | Specify the methods used to assess risk of bias in the included studies, including details of the tool(s) used, how many reviewers assessed each study and whether they worked independently, and if applicable, details of automation tools used in the process. | Methods: Data Extraction |
| Effect measures | 12 | Specify for each outcome the effect measure(s) (e.g. risk ratio, mean difference) used in the synthesis or presentation of results. | Methods: Cross-Dataset Differential Expression Meta-Analysis (Method A) |
| Synthesis methods | 13a | Describe the processes used to decide which studies were eligible for each synthesis (e.g. tabulating the study intervention characteristics and comparing against the planned groups for each synthesis (item #5)). | Methods: Study Selection |
| | 13b | Describe any methods required to prepare the data for presentation or synthesis, such as handling of missing summary statistics, or data conversions. | Methods |
| | 13c | Describe any methods used to tabulate or visually display results of individual studies and syntheses. | N/A |
| | 13d | Describe any methods used to synthesize results and provide a rationale for the choice(s). If meta-analysis was performed, describe the model(s), method(s) to identify the presence and extent of statistical heterogeneity, and software package(s) used. | Methods: Meta-Analysis, Cross-Dataset Differential Expression Meta-Analysis |

| Section and Topic | Item # | Checklist item | Location where item is reported |
|---|---|---|---|
| | | | (Method A) |
| | 13e | Describe any methods used to explore possible causes of heterogeneity among study results (e.g. subgroup analysis, meta-regression). | Methods: Differential Expression Overlap Analysis Across Datasets (Method B) |
| | 13f | Describe any sensitivity analyses conducted to assess robustness of the synthesized results. | N/A |
| Reporting bias assessment | 14 | Describe any methods used to assess risk of bias due to missing results in a synthesis (arising from reporting biases). | Methods: Data extraction |
| Certainty assessment | 15 | Describe any methods used to assess certainty (or confidence) in the body of evidence for an outcome. | N/A |
| **RESULTS** | | | |
| Study selection | 16a | Describe the results of the search and selection process, from the number of records identified in the search to the number of studies included in the review, ideally using a flow diagram. | Results: Dataset Selection, Figure 2 |
| | 16b | Cite studies that might appear to meet the inclusion criteria, but which were excluded, and explain why they were excluded. | Results: Dataset Selection |
| Study characteristics | 17 | Cite each included study and present its characteristics. | Results: Study Characteristics, Table 2 |
| Risk of bias in studies | 18 | Present assessments of risk of bias for each included study. | Results: Quality and Reproducibility Assessment |
| Results of individual studies | 19 | For all outcomes, present, for each study: (a) summary statistics for each group (where appropriate) and (b) an effect estimate and its precision (e.g. confidence/credible interval), ideally using structured tables or plots. | Results: Effect-size Based Meta-analysis (Method A) Identifies Core FRDA Genes with Substantial Cross-study Heterogeneity, Figure 5 |
| Results of syntheses | 20a | For each synthesis, briefly summarise the characteristics and risk of bias among contributing studies. | Results: Quality and Reproducibility Assessment |
| | 20b | Present results of all statistical syntheses conducted. If meta-analysis was done, present for each the summary estimate and its precision (e.g. confidence/credible interval) and measures of statistical heterogeneity. If comparing groups, describe the direction of the effect. | Results: Effect-size Based Meta-analysis (Method A) Identifies Core FRDA Genes with Substantial Cross-study Heterogeneity |
| | 20c | Present results of all investigations of possible causes of heterogeneity among study results. | Results: Recurrent Upregulated Genes Across FRDA Datasets, Recurrent Downregulated Genes Across FRDA Datasets |
| | 20d | Present results of all sensitivity analyses conducted to assess the robustness of the synthesized results. | N/A |
| Reporting biases | 21 | Present assessments of risk of bias due to missing results (arising from reporting biases) for each synthesis assessed. | N/A |
| Certainty of evidence | 22 | Present assessments of certainty (or confidence) in the body of evidence for each outcome assessed. | N/A |
| **DISCUSSION** | | | |
| Discussion | 23a | Provide a general interpretation of the results in the context of other | Discussion p2 - |

| Section and Topic | Item # | Checklist item | Location where item is reported |
|---|---|---|---|
| | | evidence. | 14 |
| | 23b | Discuss any limitations of the evidence included in the review. | Discussion p15 - 19 |
| | 23c | Discuss any limitations of the review processes used. | Discussion p18, 19 |
| | 23d | Discuss implications of the results for practice, policy, and future research. | Discussion p7 - 19 |
| **OTHER INFORMATION** | | | |
| Registration and protocol | 24a | Provide registration information for the review, including register name and registration number, or state that the review was not registered. | N/A |
| | 24b | Indicate where the review protocol can be accessed, or state that a protocol was not prepared. | N/A |
| | 24c | Describe and explain any amendments to information provided at registration or in the protocol. | N/A |
| Support | 25 | Describe sources of financial or non-financial support for the review, and the role of the funders or sponsors in the review. | Declarations |
| Competing interests | 26 | Declare any competing interests of review authors. | Declarations |
| Availability of data, code and other materials | 27 | Report which of the following are publicly available and where they can be found: template data collection forms; data extracted from included studies; data used for all analyses; analytic code; any other materials used in the review. | Declarations |

*From:* Page MJ, McKenzie JE, Bossuyt PM, Boutron I, Hoffmann TC, Mulrow CD, et al. The PRISMA 2020 statement: an updated guideline for reporting systematic reviews. BMJ 2021;372:n71. doi: 10.1136/bmj.n71. This work is licensed under CC BY 4.0. To view a copy of this license, visit https://creativecommons.org/licenses/by/4.0/

Supplementary Table 2: Systematic Review Search Strategy

| Database | Date | Results | Search Strategy |
|---|---|---|---|
| SCOPUS | 05/09/2025 | 79 | (TITLE-ABS-KEY("Friedreich's ataxia") OR TITLE-ABS-KEY("Friedreich ataxia") OR TITLE-ABS-KEY(FRDA) AND TITLE-ABS-KEY(RNA-Seq) OR TITLE-ABS-KEY(RNAseq) OR TITLE-ABS-KEY("RNA seq") OR TITLE-ABS-KEY("RNA sequencing") OR TITLE-ABS-KEY("RNA-sequencing") OR TITLE-ABS-KEY(transcriptom*)) |
| PubMed | 05/09/2025 | 57 | ("Friedreich's ataxia" OR "Friedreich ataxia" OR FRDA) AND ("RNA-Seq" OR RNAseq OR "RNA seq" OR "RNA sequencing" OR "RNA-sequencing" OR transcriptom*) |
| Gene Expression Omnibus (GEO) | 05/09/2025 | 7 | ("Friedreich's ataxia") AND "Homo sapiens"[porgn] AND ("Expression profiling by high throughput sequencing"[Filter]) |
| BioRxiv | 05/09/2025 | 40 | ("Friedreich's ataxia" OR "Friedreich ataxia" OR FRDA) AND (RNA-seq OR RNAseq OR transcriptome OR transcriptomics) |

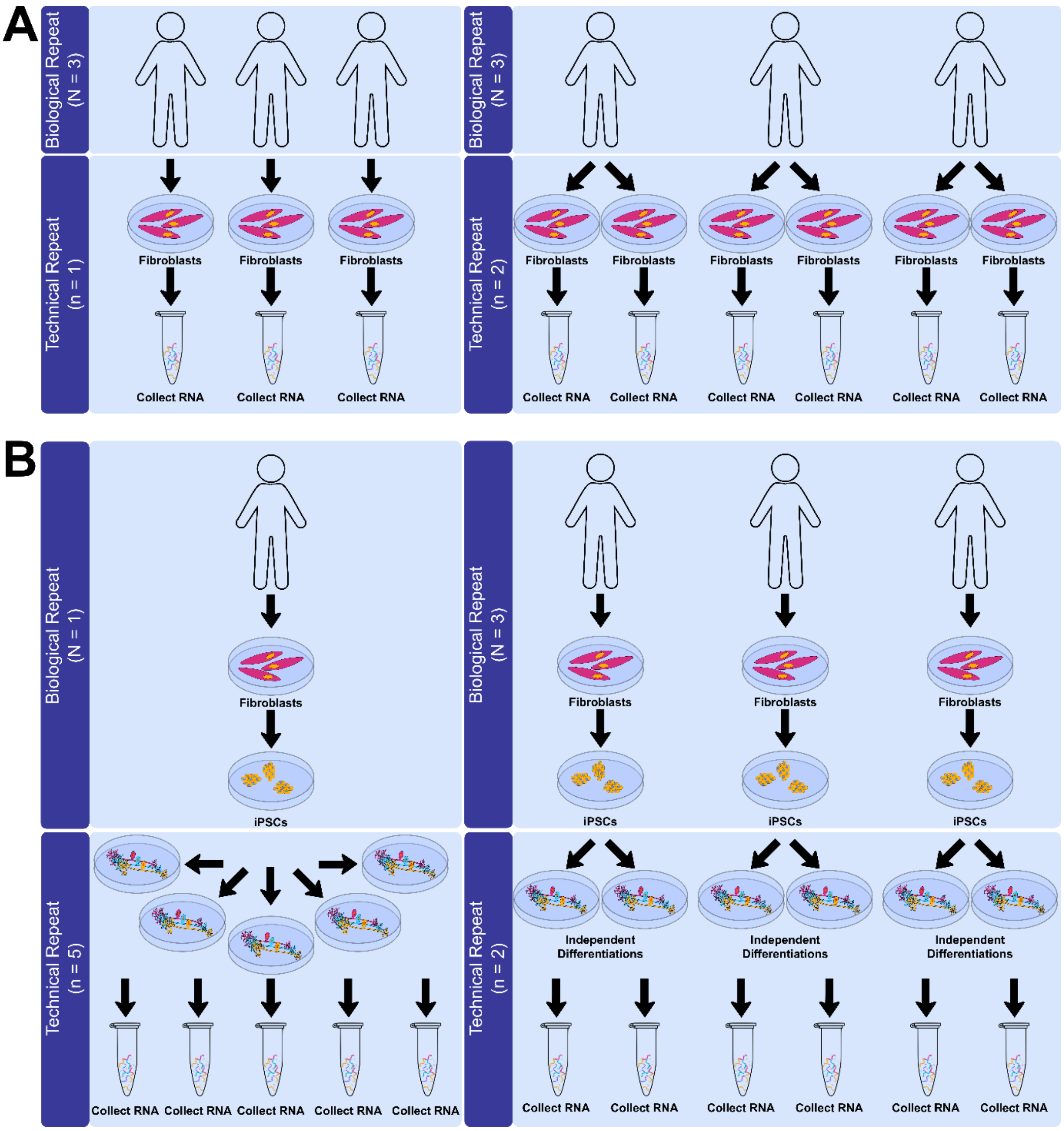


**Supplementary Figure 1:** Examples of how biological and technical replicates were defined in A) primary tissue studies and B) induced-pluripotent stem cell derived experiments for this systematic review and meta-analysis.

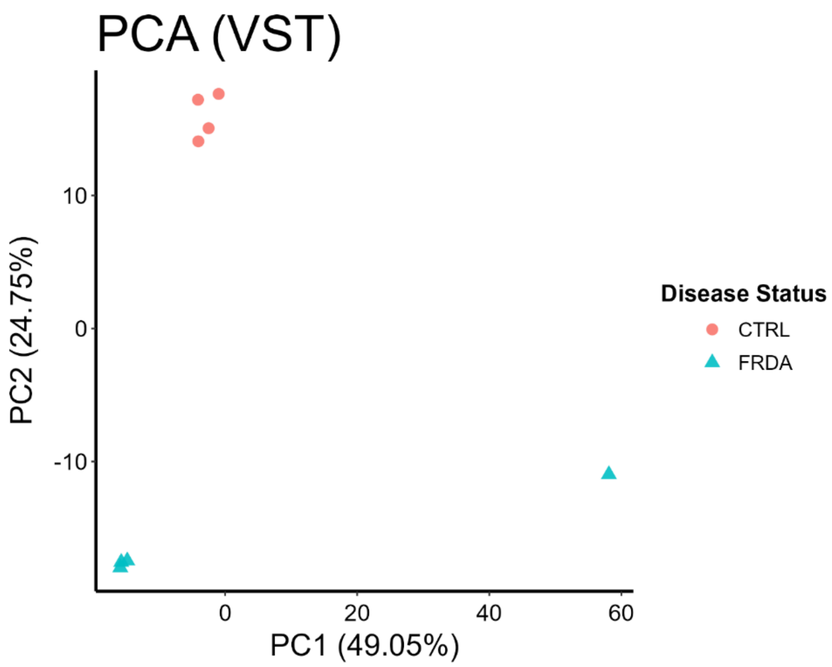


**Supplementary Figure 2:**

One FRDA sample outlier was removed from analysis within the Erwin dataset, based on principal component analysis clustering (Blue triangle at PC1 = 60%, PC2 = -10%).

**Supplementary methods 1:** Methods used to generate and characterise iPSC-derived sensory neurons, neural crest cells and lower motor neurons used in RNA-seq experiments for this study.

### iPSC Lines

RNA sequencing data generated in this study that have not been published previously were obtained from induced pluripotent stem cell (iPSC) derived neural crest cells (NCCs), iPSC-derived dorsal root ganglia sensory neurons (SNs), and iPSC-derived lower motor neurons (LMNs) from human FRDA and isogenic corrected cell lines. Two donor cell lines (FA1, FA2) and their corresponding isogenic controls (FA1ic, FA2ic), each containing a doxycycline-inducible NGN2 cassette, were used to generate NCCs ($FA1^{NCC}$, $FA2^{NCC}$, $FA1ic^{NCC}$, $FA2ic^{NCC}$) and SNs ($FA1^{SN}$, $FA2^{SN}$, $FA1ic^{SN}$, $FA2ic^{SN}$). These cell lines have been previously characterised and shown to have decreased *FXN* relative to the isogenic controls (45); however, the FA1 NGN2 line used here was derived from an independent clone (F5) that had a homozygous insert (See Supplementary Figure 3B). The GAA expansion number for each cell line is given in Supplementary Table 3. Two additional cell lines were developed from the parental FA2 and FA2ic iPSCs to drive LMN differentiation by including a doxycycline-inducible NGN2, ISL1 and LHX3 expression cassette ($FA2^{LMN}$, $FA2ic^{LMN}$) for this study. This research is covered by Human Ethics approved by the University of Wollongong Human Research Ethics Committee (HREC 202/451) and Biosafety Approval (GT19/08).

Supplementary Table 3: iPSC cell lines used to generate neural crest cells, sensory neurons and lower motor neurons.

| Cell Line | Disease Status | Sex | GAA1/GAA2 |
|---|---|---|---|
| FA1 | FRDA | Male | 867/867 |
| FA1ic | Isogenic control | Male | 0/0 |
| FA2 | FRDA | Male | 550/830 |
| FA2ic | Isogenic control | Male | 0/0 |

### Genetic Modification of iPSCs with a doxycycline-inducible NGN2, ISL1 and LHX3 expression cassette

Prior to gene editing, iPSCs were dissociated with Accutase and seeded at a density of 0.03 × $10^6$ cells/$cm^2$ in Essential 8 (E8) medium supplemented with the ROCK inhibitor Y-27632 (10 µM). CRISPR/Cas9 reagents targeting the CLYBL safe-harbour locus were delivered as a ribonucleoprotein (RNP) complex composed of a crRNA:tracrRNA duplex and high-fidelity Cas9 nuclease (IDT). 1 × $10^6$ cells were transfected with the RNP complex together with 2 µg HDR donor plasmid (Addgene #124230; CLYBL-TO-hNIL-BSDmApple) and 0.4 µg Addgene plasmid #41856, which transiently expresses mouse p53DD to enhance homology-directed repair. The construct is given in Figure 3A. Transfections were performed using Lipofectamine Stem Transfection Reagent (Invitrogen). Forty-eight hours post-transfection, cells were subjected to blasticidin selection (10 µg/mL) to enrich for cassette-integrated clones. Single-cell–derived colonies were subsequently obtained by limiting dilution.

Targeted integration of the gene cassette at the CLYBL locus was verified by junction PCR, confirming homozygous insertion (Figure 3B). Genomic DNA was isolated using the PureLink Genomic DNA Mini Kit (Thermo Fisher Scientific). PCR amplification was performed with Invitrogen Platinum Green Hot Start PCR 2× Master Mix using primers listed in Supplementary Table 4, and reactions were run on an Eppendorf Mastercycler Pro.

Supplementary Table 4: Primers

| | **Target** | **Forward/Reverse primer (5'-3')** |
|---|---|---|
| Genotyping primers | FW WT primer | TGACTAAACACTGTGCCCCA |
| | RV WT primer | AGGCAGGATGAATTGGTGGA |
| | FW Insert primer | CAGACAAGTCAGTAGGGCCA |
| | RV insert primer | AGAAGACTTCCTCTGCCCTC |
| sgRNA/crRNA sequence including PAM sequence (bold) | CLYBL CRISPR sgRNA | ATGTTGGAAGGATGAGGAAA **TGG** |

NGN2, ISLET1 and LHX3 overexpression was verified after 48 h of doxycycline addition in the iPSCs by immunostaining (Supplementary Figure 3C). Both cell lines remained pluripotent after genetic modification, expressing the pluripotency markers OCT4 and SOX2 (Supplementary Figure 3D). Karyotyping with the CytoScan Opima Assay showed a 4,926 kbp gain on chromosome x in cytoband q25-q26.2 in the FA2$^{LMN}$ cell line. Finally, *FXN* was shown to be significantly decreased by 87.9 % in the FRDA FA2$^{LMN}$ iPSCs compared to the FA2ic$^{LMN}$ isogenic controls ($t(4.07) = 7.20$, $p = 0.0019$) (Supplementary Figure 3E).

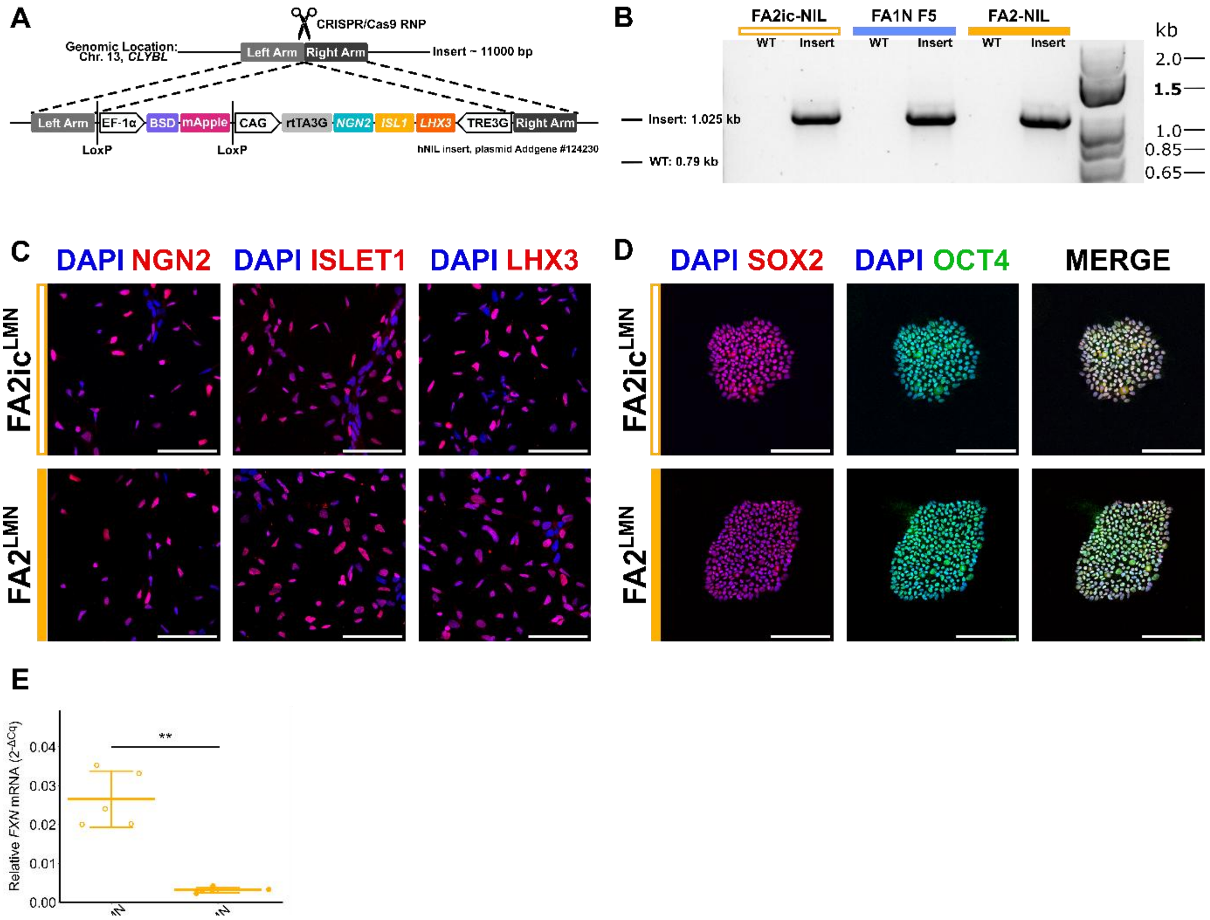


Supplementary Figure 3: Genetic modification and validation of inducible lower motor neuron iPSC lines. A) Doxycycline-inducible gene construct used in iPSC cell line FA2. B) Junction PCR validation confirming correct genomic integration of the inducible cassette. All cell lines were homozygous for the insertion. C) NGN2, ISLET1 and LHX3 was expressed in iPSCs after doxycycline treatment for 48 hours. D) iPSCs stained positive for pluripotency markers SOX2 and OCT4 after genetic modification. E) *FXN* gene expression was significantly decreased in the $FA^{LMN}$ vs isogenic corrected $FA2ic^{LMN}$ iPSCs (**$p$ < 0.01).

### Cell Culture

All iPSCs used have tested negative for mycoplasma and not undergone spontaneous differentiation. iPSC lines were maintained on 10 µg/mL vitronectin XF™ Matrix (StemCell Technologies) coated T-25 tissue culture flasks (Interpath) in feeder-free TeSR™-E8 culture media (StemCell Technologies) at 37 °C, and 5 % carbon dioxide ($CO_2$) until 60-80 % confluency was reached. For continuous proliferation, iPSCs were passaged into a new vitronectin-coated T-25 flask by washing the cells with DPBS- (ThermoFisher), followed by 3-minute incubation in 0.5 mM ethylenediaminetetraacetic acid (EDTA) (ThermoFisher Scientific). The EDTA was then removed, and cells resuspended in 2 mL of TeSR-E8 media by tapping the flask. iPSCs were then transferred to a new T25 flask at a 1:2 – 1:10 dilution depending on the desired cell density for continued maintenance.

### iPSC differentiation to neural crest cells

Neural Crest Cells (NCCs) were generated from iPSC cell lines with the *NGN2* insert using the STEMDiff Neural Crest Kit (Stem Cell Technologies, #08610). When iPSCs reached approximately 70 % confluency, colonies were lifted with accutase (7 minutes, 37 °C, 5 % $CO_2$) and plated as single cells on Matrigel-coated 6 well plate at a density of 400,000 – 700-000 cells per well. Daily media changes following manufacturer instructions were performed for 7 days. On day 7 NCC differentiation was complete, and cells were collected for RNA, plated on coverslips for immunostaining and further differentiated into sensory neurons. NCC identity was confirmed through SOX10 and p75NTR staining (Supplementary Figure 4).

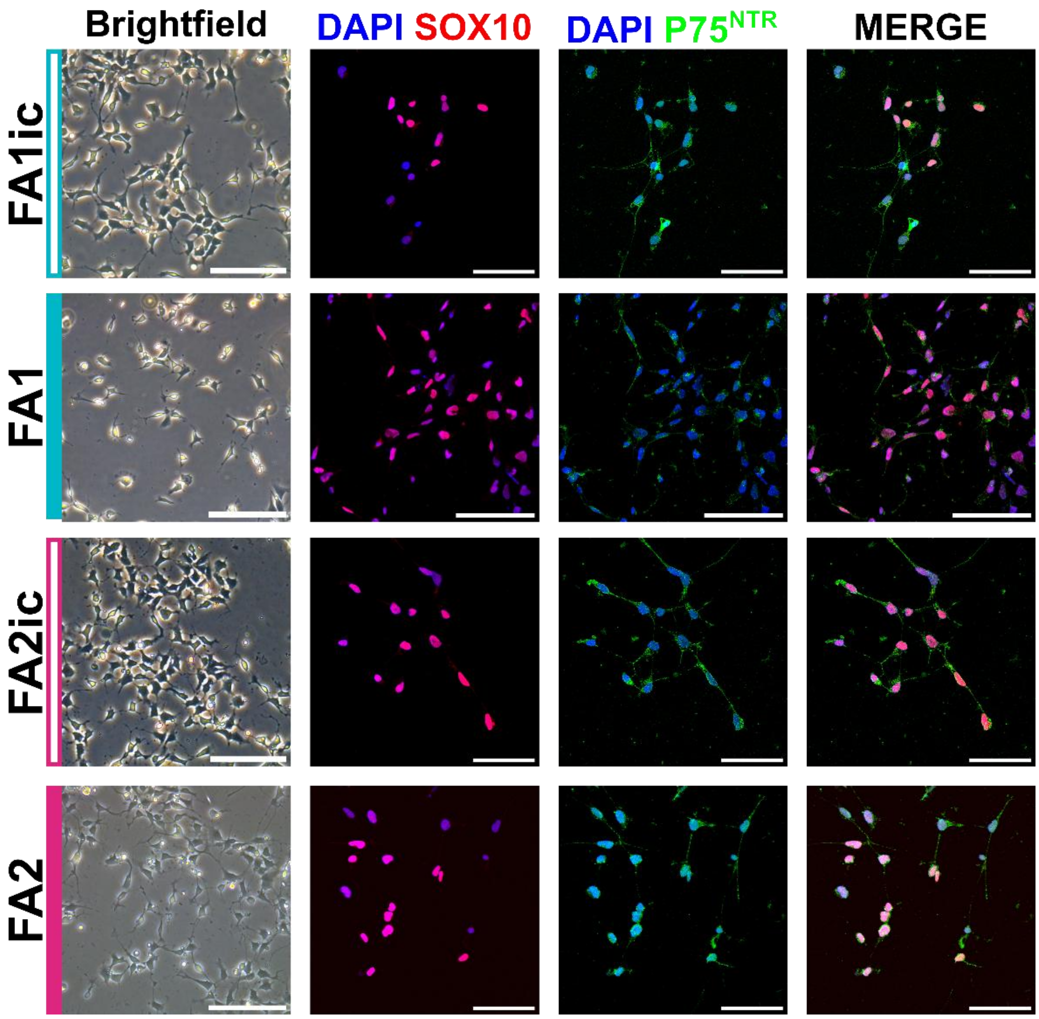


Supplementary Figure 4: All iPSC-derived neural crest cells displayed typical neural crest cell morphology expressed neural crest cell markers SOX10 and p75-NTR.

**iPSC differentiation to sensory neurons**

Sensory neurons were differentiated from NCCs using a combined small-molecule and transcription factor protocol. Tissue culture plates were first coated with 0.05 mg/mL Poly-D-Lysine for 1 hour at room temperature, followed by coating with Matrigel overnight at 4 °C. NCCs were seeded at a density of 1.7 x $10^4$ cells/$cm^2$. Sensory neurons were induced by overexpression of NGN2 with 2 µg/mL doxycycline for the first 120 hours post-plating. Media was changed every 48 – 72 hours. Cells were maintained in neurobasal media supplemented with 1X N-2, 1X B-27 (without vitamin A), 1X insulin transferrin selenium A, 2 mM GlutaMAX™ (all from Gibco™), for the first five days of differentiation. On day six, cells were transitioned into BrainPhys neuronal medium with 1X NeuroCult SM1 without vitamin A, 0.5X N2 Supplement-A (Stem Cell Technologies) using a 25:75, 50:50, 75:25, 100:0 ratio of BrainPhys™ : neural media with each media change. From day 14, BrainPhys™ media was changed until mature sensory neurons were present at 21 days of differentiation. All media changes included 10 µM Y-27632, 10 ng/mL GDNF, 10 ng/mL BDNF, 10 ng/mL NT-3 and 10 ng/mL β-NGF (all from Stem Cell Technologies) throughout differentiation. Cytosine β-D-arabinofuranoside (AraC; 2.5 µM, Sigma-Aldrich) was applied for 48 h between days 8 and 10 to inhibit proliferating cells. At day 21, sensory neurons were either collected for RNA or fixed for immunostaining. All iPSC-derived sensory neurons were confirmed to express the sensory neuron markers NF200, BRN3A and ISLET1 (Supplementary Figure 5A).

**iPSC differentiation to lower motor neurons**

The iPSCs with a stably integrated human NIL under a tetracycline-inducible promoter were differentiated to LMNs as following: Briefly, on day 0 the iPSCs were single-cell dissociated using Accutase (Thermo Fisher) and plated in a Matrigel-coated (in vitro technologies) dish in E8 medium (Stemcell Technologies) and 10 mM Rock inhibitor Y-27632 (Stemcell technologies). On day 1, the media was changed to NIM:DMEM/F12, 1X N2, 1X Non-Essential Amino Acids, 1X Glutamax (all reagents were from Thermo Fisher), containing 10 mM Rock inhibitor Y-27632, 2 mg/mL doxycycline (Sigma) and 0.2 mM Compound E (Stem Cell Technologies). On day 3, 0.3x$10^6$ cells were replated per one well of a poly-D-lysine (Thermofisher) and Matrigel-coated 12-well plate in NIM with 10mM Rock inhibitor Y-27632, 2 mg/mL doxycycline (Sigma-Aldrich), 0.2 mM Compound E (Stem Cell Technologies), 1 mg/mL Laminin (Thermofisher) and 40 mM BrdU (Sigma-Aldrich). On day 4, the media was changed to Neurobasal media with 1X B27 Supplement (Thermo Fisher), 1x N2 (Thermofisher), 2 mg/mL doxycycline,1mg/mL Laminin, 20 ng/mL BDNF (Stemcell technologies), 20 ng/mL GDNF (Stemcell technologies) and 10 ng/mL NT3 (Stemcell technologies). Media changes were performed every 2-3 days. From Day 10 onwards cells were transitioned into BrainPhys neuronal medium with 1X NeuroCult SM1, 0.5X N2 Supplement-A (Stem Cell Technologies) using a 25:75, 50:50, 75:25, 100:0 ratio of BrainPhys™: neurobasal media with each media change, omitting doxycycline from this day onwards. iPSC-derived LMNs expressed the LMN markers HB9, CHAT, ISLET1 and SMI32 (Supplementary Figure 5B). *FXN* was significantly reduced in each FRDA versus isogenic control comparison ($p < 0.05$).

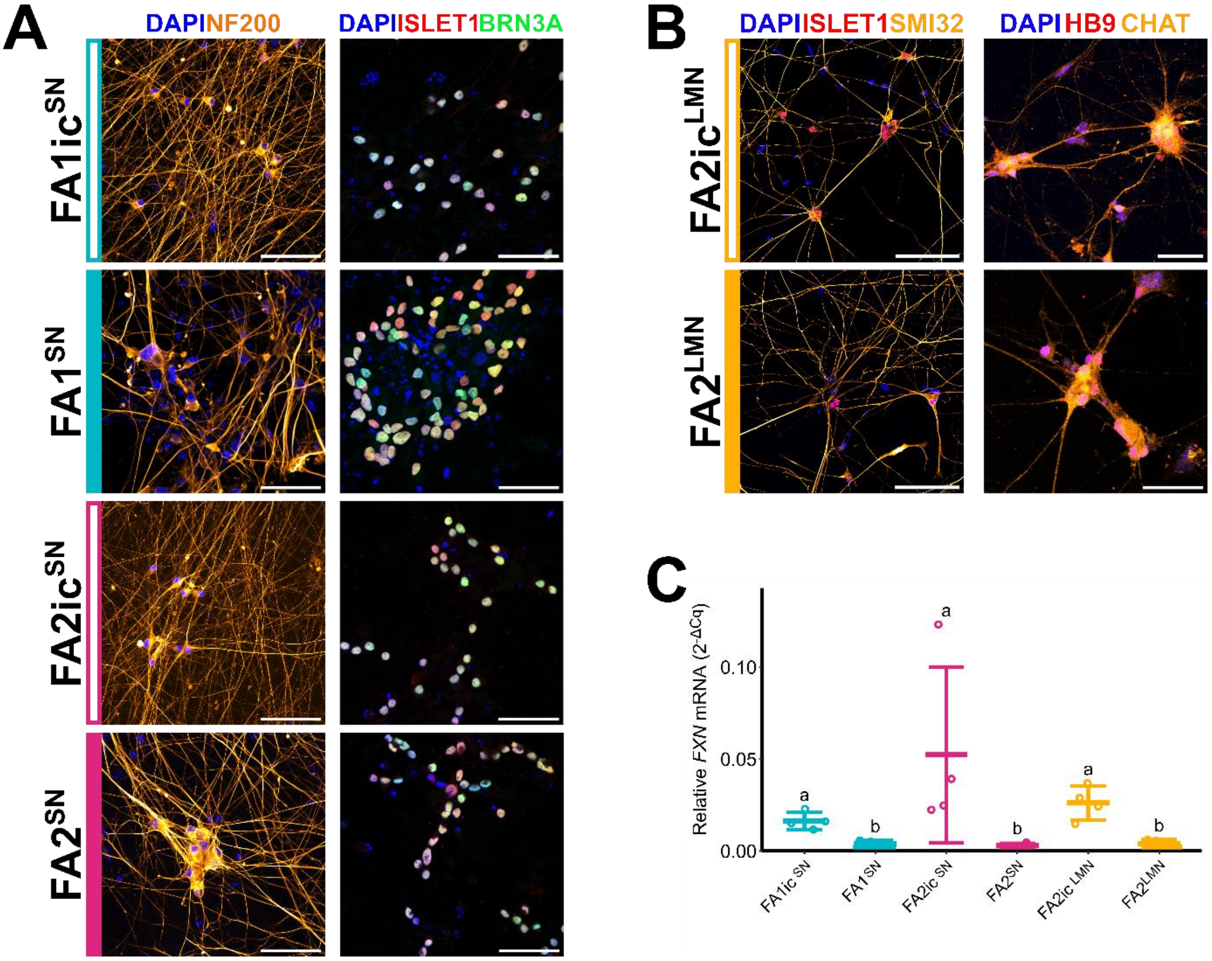


Supplementary Figure 5: Characterisation of iPSC-derived sensory neurons and lower motor neurons used in RNA-seq experiments. A) All FRDA ($FA1^{SN}$, $FA2^{SN}$) and isogenic corrected ($FA1ic^{SN}$, $FA2ic^{SN}$) sensory neurons exhibited typical morphology and expressed sensory neuron markers NF200, BRN3A and ISLET1. B) iPSC-derived lower motor neurons from FRDA ($FA^{LMN}$) and isogenic corrected ($FA2ic^{LMN}$) cell lines expressed canonical lower motor neuron markers ISLET1, SMI32, HB9 and CHAT. C) *FXN* expression measured by RT-qPCR was significantly reduced in each FRDA versus control comparison. Groups with the same letter are not significantly different from each other.

**RNA Extraction**

To extract total cellular RNA for RT-qPCR and RNA sequencing, the PureLink RNA Mini Kit (ThermoFisher Scientific) was implemented according to the manufacturer's instructions, with modifications. Inside a fume hood, cell cultures were washed twice with cold phosphate buffered saline (PBS) and lysed using the kit's lysis buffer supplemented with 1:100 β-mercaptoethanol. Sensory and lower motor neuron cultures formed cohesive sheets that were selectively detached to enrich for neuronal populations, while non-neuronal contaminants remained adherent. Equal parts of 70% ultra-pure ethanol (E7023, Sigma) were added to the lysate to precipitate the RNA. This was then placed into a PureLink RNA spin column. To purify the RNA, multiple washing and centrifugation steps at 13,000 x g were performed. Briefly, 700 µL of wash buffer I was added to the column and centrifuged for 1 minute. Next, 500 uL of wash buffer II was added and centrifuged (1 minute). Wash buffer II was added again to purify the RNA further, and centrifuged for 2 minutes. This was followed by an additional centrifugation step for 3 minutes. Finally, 30 µL of UltraPure™ DNase/RNase-Free distilled water (10977015, ThermoFisher) was added to the column and centrifuged for 2 minutes. The yield and purity of RNA were assessed using a NanoDrop 2000c UV-Vis spectrophotometer (ThermoFisher Scientific).

For RT-qPCR experiments, cDNA synthesis of RNA samples was performed according to the iScript™ gDNA Clear cDNA Synthesis Kit (1725035, Bio-Rad). RT-qPCR was used to determine *FXN* expression. Target gene transcripts were amplified in a QuantStudio™ 5 Real-Time PCR system in a 0.1 mL 96-well PCR plate (Applied Biosystems) using TaqMan probes (ThermoFisher). The PCR reaction mix consisted of 1X (5 µL/reaction) TaqMan™ Fast Advanced Master Mix (Applied Biosystems), 1X (0.5 µL/reaction) TaqMan™ frataxin (Hs00175940_m1), GAPDH (Hs02786624_g1) or TATA box binding protein (TBP; Hs00427620_m1) probes and 20 ng/µL (2 µL/reaction) sample cDNA with 2.5 µL of UltraPure water for a final volume of 10 µL. The GAPDH and TBP probes were used as the housekeeping reference genes. The RT-qPCR reaction conditions were 2 minutes at 50°C, followed by 20 seconds at 95°C to activate the polymerase, then 40 cycles of 1 second at 95°C and 20 seconds at 60°C. Each target was performed in triplicate. Outlier replicates were excluded if the standard deviation of the three replicates' quantification cycle (Cq) value was > 0.5. Each target had an additional non-template control with no cDNA and was run in triplicate. Expression analyses were conducted using QuantStudio™ Design and Analysis Software (v2.6.0) (Applied Biosystems), and quantified using the ProntoPCR analysis software (https://marniemaddock.github.io/ProntoPCR/). Expression is given as relative mRNA ($2^{-\Delta Cq}$).

**Immunostaining**

iPSCs, NCCs, SNs and LMNs cultured on 13 mm glass coverslips were fixed with 4 % w/v paraformaldehyde in phosphate-buffered saline (PBS) for 20 minutes at room temperature. Fixed cells were then washed three times in PBS, permeabilised with 0.3 % Triton X-100 buffer (for iPSCs, NCCs) or 0.1 % Triton X-100 buffer (for SNs and LMNs) for 10 minutes, blocked in 10 % v/v normal donkey serum in PBS for 1 hour. Samples were then incubated overnight at 4 °C with primary antibodies (Table X) prepared in 10 % v/v normal donkey serum. Then, samples were washed 3 times in 0.1 % Triton-X-100 Buffer, incubated with secondary antibodies (Supplementary Table 5) in 10 % v/v normal donkey serum for 2 hours at room temperature and mounted on glass slides using ProLong Glass antifade mountant (Invitrogen). All slides were imaged using a Leica SP8 confocal microscope using a 40x or 63x oil-immersion objective.

Supplementary Table 5: Antibodies used in characterisation of iPSCs, and derived neural crest cell, sensory neuron and lower motor neuron cell types.

| **Primary Antibodies** | **Dilution** | **Catalogue Number, Supplier, RRID** |
|---|---|---|
| Mouse anti-OCT4 | 1:200 | SC-5279, Santa Cruz Biotechnology, RRID:AB_628051 |
| Goat anti-SOX2 | 1:200 | AF2018, R&D Systems, RRID:AB_355110 |
| Rabbi anti-NGN2 | 1:500 | PA5-78556, Thermofisher, RRID:AB_2736211 |
| Rabbit anti-ISLET1 | 1:500 | Ab20670, Abcam, RRID:AB_881306 |
| Rabbit anti-LHX3 | 1:200 | PA1-29491, Thermofisher, RRID:AB_2135675 |
| Goat anti-SOX10 | 1:200 | AF2864, R&D Systems, RRID:AB_442208 |
| Mouse anti-p75NTR | 1:500 | M-1818-100, Biosensis |
| Mouse anti-BRN3A | 1:500 | MAB1585, Millipore, RRID:AB_94166 |
| Mouse anti-NF200 | 1:1000 | N0142, Sigma-Aldrich, RRID:AB_477257 |
| Mouse anti-SMI32 | 1:500 | Ab7795, Abcam, RRID:AB_206084 |
| Rabbit anti-HB9 | 1:250 | PA5-23407, Thermofisher, RRID:AB_2540929 |
| Mouse anti-ChAT | 1:200 | MA5-31382, Thermofisher, RRID:AB_2787019 |
| Donkey Anti-Mouse IgG H&L (Alexa Fluor 488) | 1:500 | Ab150109, Abcam, RRID:AB_2571721 |
| Donkey Anti-goat IgG H&L (Alexa Fluor 647) | 1:500 | Ab150135, Abcam, RRID:AB_2687955 |
| Donkey Anti-Rabbit IgG H&L (Alexa Fluor 555) | 1:500 | Ab150062, Abcam, RRID:AB_2801638 |

**RNA sequencing library preparation**

Total RNA was extracted from NCCs, SNs and LMNs using the PureLink™ RNA Mini kit (ThermoFisher) and submitted to the Ramaciotti Centre for Genomics (UNSW Sydney, Australia) for quality control, library preparation and sequencing. All samples had a high RNA integrity (RIN $\geq$ 9) as measured on an Agilent TapeStation. rRNA depleted libraries were prepared using the Illumina Stranded Total RNA Ribo-Zero Plus kit and sequenced on the Illumina NovaSeq X Plus platform to generate 100 bp paired-end reads. Sequencing yielded between 76.8 – 184.8 million read pairs per sample.

**Supplementary Methods 2: Candidate Biomarker Panel Identification Extended Methods**

Based on the most consistently up or downregulated genes in FRDA, we defined a candidate transcriptomic biomarker panel comprising the top 40 ranked genes. To identify cell-type specific biomarkers and pathways, analyses were additionally performed separately for major cell types, including fibroblasts (Napierala, Vilema-Enriquez, Wang), lymphoblastoid cells (Erwin, Chutake), cardiomyocytes (Lees, Li), sensory neurons (Lai - PNS, Maddock - SN), CNS/LMNs (Lai - CNS, Maddock - LMN) and all neurons (Lai, Maddock, Mishra). The top 10 up and downregulated genes based on rankings are displayed for visualisation purposes. These ranked lists were based on the number of studies in which a gene was significantly (FDR < 0.05; FRDA vs control) upregulated ($n_{up}$) or downregulated ($n_{down}$). Directional consistency was calculated using a direction score for each gene as follows:

$$Direction\ score = \frac{n_{up} - n_{down}}{n_{up} + n_{down}}$$

The score ranges from -1 (consistently downregulated) to +1 (consistently upregulated) with values near zero indicating weak or inconsistent directionality. Effect size magnitude was summarised using the median $\log_2$ fold-change across all studies. Genes were prioritised using a hierarchical ranking method that favoured genes with (i) the highest number of studies showing significant up or downregulation overall, (ii) the fewest conflicting studies in terms of significant up versus downregulation, iii) stronger directional agreement as quantified by the direction score, and (iv) larger absolute median $\log_2$ fold-change as a final tie-breaker. The top 40 genes were then evaluated in an independent peripheral blood microarray dataset from 418 individuals with FRDA and 93 unaffected controls to assess their potential relevance as blood-based biomarkers (50).
